# A flexible workflow for building spectral libraries from narrow window data independent acquisition mass spectrometry data

**DOI:** 10.1101/2021.11.22.469568

**Authors:** Lilian R. Heil, William E. Fondrie, Christopher D. McGann, Alexander J. Federation, William S. Noble, Michael J. MacCoss, Uri Keich

## Abstract

Advances in library-based methods for peptide detection from data independent acquisition (DIA) mass spectrometry have made it possible to detect and quantify tens of thousands of peptides in a single mass spectrometry run. However, many of these methods rely on a comprehensive, high quality spectral library containing information about the expected retention time and fragmentation patterns of peptides in the sample. Empirical spectral libraries are often generated through data-dependent acquisition and may suffer from biases as a result. Spectral libraries can be generated *in silico* but these models are not trained to handle all possible post-translational modifications. Here, we propose a false discovery rate controlled spectrum-centric search workflow to generate spectral libraries directly from gas-phase fractionated DIA tandem mass spectrometry data. We demonstrate that this strategy is able to detect phosphorylated peptides and can be used to generate a spectral library for accurate peptide detection and quantitation in wide window DIA data. We compare the results of this search workflow to other library-free approaches and demonstrate that our search is competitive in terms of accuracy and sensitivity. These results demonstrate that the proposed workflow has the capacity to generate spectral libraries while avoiding the limitations of other methods.

## 1 Introduction

Data independent acquisition (DIA) mass spectrometry (MS) has emerged as a powerful method to simultaneously quantify tens of thousands of peptides in a single MS run.^1;2^ Unlike data dependent acquisition (DDA) tandem mass spectrometry methods which select precursor ions to fragment in real time, in DIA the instrument cycles through a series of pre-defined isolation windows to fragment all ions in a specific mass-to-charge (*m/z*) range.^3^ Decreasing the precursor isolation width comes at a cost to either total mass range covered or the number of points across the peak, and therefore may not be a viable option. Consequently, DIA tandem mass spectra may contain ions drawn from mass ranges of 20 Da or more and can be highly complex due to the presence of product ions from multiple co-isolated peptides. As DIA rises in popularity, the increasing volume and complexity of data presents computational challenges for peptide detection.

Due to the massive amounts of data generated by modern mass spectrometers, database searches are essential to determine which precursor peptide generated a spectrum at a large scale. Traditional search methods in proteomics use a “spectrum-centric” approach^4^ to identify the peptide sequence that best explains a given tandem mass spectrum (MS2). These searches are optimized for situations where each spectrum represents exactly one peptide, an assumption which originates from typical DDA mass spectrometry data, but is often invalid for unprocessed DIA mass spectrometry data. Therefore “peptide-centric” methods were developed, which aim to detect the best evidence for each possible peptide sequences in a set of spectra. Methods such as PECAN^5^ employ a peptide-centric method to detect peptides in DIA data based on a protein sequence database. Unfortunately, the large search space limits the statistical power of these searches, and it is not possible to detect modified peptides without sacrificing statistical power due to multiple hypothesis testing.^5^ This limitation is inherent to peptide-centric searches as the candidate peptide search space increases with added search complexity, while spectrum-centric searches largely avoid this problem due to a limited number of spectra.

One DIA search strategy is to group co-eluting fragments and precursor ion signals and assign them to groups of pseudo-spectra that can be searched using spectrum-centric DDA database search methods.^6–9^ While this method enables the detection of post-translational modifications without a spectral library, it relies on the presence of a precursor MS1 signal, which may not be present for all peptides that are detectable at the product ion MS2 level. This problem is common in trapping instruments such as Orbitraps where automatic gain control modulates the fill time to maximize the number of ions in the mass analyzer while minimizing negative impacts of overcrowding such as space charging and ion coalescence.^10^ Automatic gain control enables the MS2 spectra to be substantially more sensitive than an MS1 spectrum by filling the trap with a narrow *m/z* range and excluding ions outside the window.^11^

Another way to increase statistical power of DIA searches is to use spectral libraries which contain information about the retention times and fragmentation patterns of peptides that might be in a sample. Using this information, spectral libraries can be used to perform targeted signal extraction on DIA tandem mass spectra to detect and quantify peptides with great sensitivity and precision.^1;12–16^ By incorporating additional features into scoring models, the use of libraries significantly boosts the performance of peptidecentric searches.^17^ Because analyses using spectral libraries are limited by the depth and quality of the library, an incomplete or low quality spectral library can derail analysis of even the best data.^18^ Thus, the method of library generation is of utmost importance in DIA mass spectrometry experiments.

Traditionally, spectral libraries have been generated by offline fractionation via liquid chromatography followed DDA tandem mass spectrometry. These experiments are time consuming, sample intensive, and yield libraries which may not represent the behavior of peptides in the sample used for quantitative DIA mass spectrometry analysis.^13^ The inaccuracy of DDA-based spectral libraries arises because offline fractionation changes the composition of the sample matrix relative to the unfractionated sample, causing shifts in retention time.^13;19^ Additionally, DDA methods typically use different, charge state-dependent collision energies as opposed to DIA methods that typically assume the same charge state for every fragmentation spectrum, creating differences in peptide fragmentation patterns.^13;19^ To conserve sample and time, publicly available spectral libraries generated from DDA methods can be used in place of creating a DDA fractionated library in-house. However, even high quality publicly available libraries, such as the Pan-Human library,^20^ may lack crucial peptides specific to a sample and are not available for every possible experimental condition and organism. This problem may be exacerbated when specific post-translational modifications (PTMs) are of interest that may not be detectable in a library without enrichment or may be unique to a specific set of experimental conditions.

An alternative method to overcome the problems with DDA-based spectral library generation is to use a computational model to predict entire spectral libraries. Recently, tools to predict peptide fragmentation and retention time have improved such that it is possible to accurately predict these features for all possible unmodified peptides in a proteome, producing spectral libraries that are entirely predicted *in silico.*^21–23^ Programs such as EncyclopeDIA can integrate predicted spectral libraries to detect and quantify peptides in quantitative DIA mass spectrometry data.^13;21^ However, robust methods to predict which peptides are likely to be detectable in a sample are not readily available and these searches are hindered by the large search space which imposes a limit on sensitivity.^13;24^ Additionally, these methods need to be trained specifically for individual post-translational modifications which can greatly limit their use.

To reduce the search-space and calibrate spectral libraries to a specific sample, gas-phase fractionation of a pooled sample can be used to obtain deep coverage of the whole sample space. In this method, a single sample is injected six times, and DIA mass spectrometry is performed using narrow (4-*m/z*) staggered isolation windows.^25^ This acquisition method, similar to the PAcIFIC method,^26^ can be used to generate sample-specific chromatogram libraries based on the predicted library which increase the power of the search.^13;19^ Because these searches are limited by the scope of the predictable library, which typically only covers unmodified and fully tryptic peptides, a method to generate libraries that cover more peptides is essential. Thus, limitations in spectral library generation have complicated the widespread adoption of DIA in many circumstances.

To address the current obstacles in library construction for DIA data, we propose a workflow to detect peptides directly from gas-phase fractionated, narrow-window DIA data using a modified spectrum-centric approach outlined in Supplemental Algorithm 1 (Figure 1). The method involves a spectrum-centric database search to score all possible spectrum/peptide combinations – where here each peptide entry consists of a peptide sequence with a specified charge state. Then, a series of steps for score calibration and a novel false-discovery rate estimation method are used to generate a list of peptides and their corresponding spectra. We use the results of this search to build spectral libraries that include PTMs, thereby enabling us to detect and quantify these peptides in wide-window DIA mass spectrometry data. In contrast to other spectrum-centric approaches, this search workflow utilizes information from the time domain to generate a spectral library. We demonstrate that this search method outperforms DIA-Umpire,^8^ a pseudo-spectrum based method, to detect phosphorylated peptides without the use of a spectral library. Additionally, we show that the peptides detected with our method can be accurately quantified in DIA mass spectrometry data. The DIA-based, empirically generated spectral library produced by our proposed workflow overcomes the limitations of many DIA search methods and simplifies the generation of high-quality libraries for many experiments that were previously unable to use DIA mass spectrometry.

**Figure 1:**
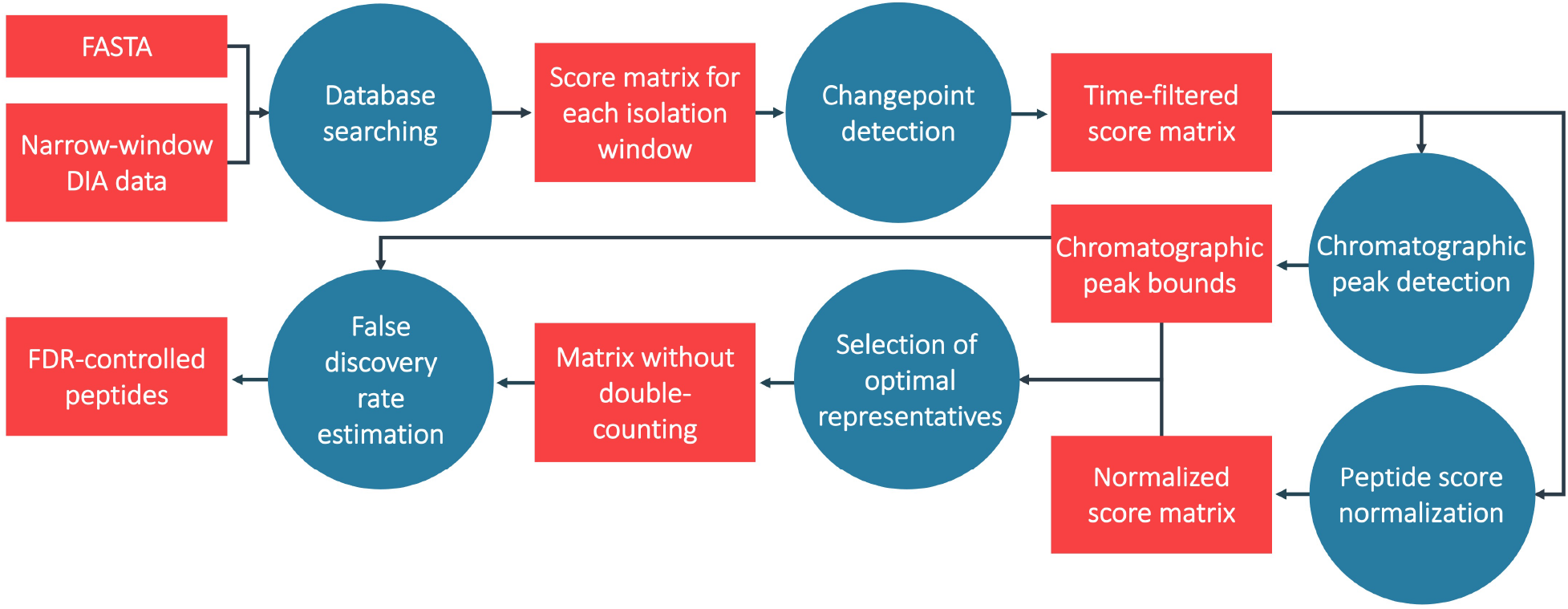
Search workflow. Workflow for direct searching of narrow-window DIA data. Raw data is searched against a FASTA file and the search results are converted into a matrix of scores. Then, spectra at the beginning and end of the run are removed from the matrix using changepoint detection, the chromatographic peak width is determined, scores are normalized, and highly correlated peptides (at their respective charge-states) are competed against one another directly. Finally, this set of peptides is subjected to FDR control where peptides are considered at multiple precursor charge-states.

## 2 Approach

Peptide detection in DIA data presents unique challenges and opportunities that should be considered when designing a search method. The proposed workflow aims to address these challenges and capitalize on the unique information provided by DIA tandem mass spectrometry data. As with any peptide database search, it is important to select a sensitive and powerful score metric to quantify the quality of each peptide-spectrum match (PSM). Additionally, traditional spectrum-centric type searches fail to utilize the time-domain information present within DIA data and may struggle to account for highly chimeric spectra. However, simply retaining the top *n* best-scoring peptide candidates per spectrum with no restrictions can lead to artificially large numbers of peptide detections because multiple peptides sequences sharing predicted fragments may score highly while only one is present. More generally, due to the complexity of DIA data, traditional target/decoy competition methods can be insufficient for DIA searches.

Here, we present a workflow to detect peptides in narrow-window DIA data that is designed to address the challenges of DIA searches with a multi-step procedure, which we describe in detail in the following sections. Note that we use the term peptide to denote a peptide sequence and charge state combination. Our notation is summarized in Supplemental Table 1 and the overall procedure is summarized in (Figure 1).

### 2.1 Database searching

Our procedure begins by conducting a Tide search^27^ using the XCorr score function^28^ with Tailor calibration^29^ and a sufficiently large top-match parameter so that every possible mass spectrum/peptide combination is assigned a score. The Tide search results are stored in matrices, each covering a single *2-m/z* precursor isolation window, where each row represents a theoretical peptide at a specific charge state, each column represents a tandem mass spectrum, and each entry is the Tailor calibrated XCorr score of the match between the corresponding peptide and spectrum (Supplemental Figure 1, Supplemental Algorithm 2). Each matrix *M* consists of *m* peptides *P* and *n* mass spectra *S*, where *P* and *S* are specific to each 2-*m/z* isolation window, and it contains no missing values. In a typical experiment, there are 50 matrices for each of the six gas phase fractions. For a 90-minute gradient searched against all unmodified fully tryptic yeast peptides, each of these 300 matrices is approximately 2000 peptides by 1500 spectra.

### 2.2 Changepoint detection

DIA data includes uninteresting mass spectra at the beginning and end of the run before and after peptides are eluting. The matrix columns corresponding to these mass spectra have score distributions that are distinct from the rest of the run, and are identified using changepoint detection (Supplemental Algorithm 3). Specifically, we considered (in this order): two, three, or four changepoints in the mean and variance of the median of the score distribution for each spectrum. The changepoints were identified using binary segmentation and if the number of spectra between the first and last changepoints was at least 50% of the total number of spectra, then we stopped at that number of changepoints and continued the rest of the analysis using only the spectra between those changepoints (Supplemental Figure 2).^30–32^ This process focuses our analysis on mass spectra where peptides are likely to be present, thus ensuring more accurate score calibration.

### 2.3 Chromatographic peak detection

As peptides elute from the column, we expect to observe them in multiple consecutive mass spectra, a feature that could potentially help to distinguish between true and false detections, but using this information is not straightforward. Notably, the associated chromatographic peak widths may vary due to many factors. Therefore, we propose a computational method to deduce the chromatographic peak width of each peptide.

In the case of DIA spectra, determining the chromatographic peak boundaries is equivalent to assigning each peptide *p_i_* a time-consecutive sequence of MS2 spectra which contain fragment ions that can be ex-plained by *p_i_*. Computationally this amounts to scanning the *i^th^* row of the PSM matrix *M* for a stretch of column-consecutive high scoring entries anchored at the maximal entry for that row. While this process is conceptually straightforward, the baseline score distributions vary substantially between peptides. Therefore, we found that further normalization applied to each row significantly improves our ability to assign correct peak widths. Specifically, we use robust z-scores that provide normalization that is robust to outliers that appear when peptides have a few high scoring spectra.

Hence, matching the consecutive sequence of MS2 spectra to a peptide to assign peak width amounts to applying row-wise robust z-score normalization, and then identifying the maximal entry for each row (peptide). Then, we extend the sequence by adding as many entries to the left and right of the maximal entry with a score ≥ 70% of the maximal entry. When executing target/decoy competition-based FDR control, it is essential that each target/decoy peptide pair shares the same width group. Therefore, we assign each target/decoy pair the maximum of the two assigned widths. In Supplemental Algorithm 4 we provide more details on the approach.

Following this last step, we compete each target peptide with its corresponding decoy so that only the larger scoring one is retained while the other is removed from consideration (Supplemental Algorithm 6). This direct competition is essential to our false-discovery rate control and it is performed early in the procedure here to reduce the computational overhead of subsequent steps.

Even with accurate chromatographic peak width boundaries, it is not obvious how to use this information. After exploring several options, including methods to reward peptides with wider peaks, we ended up exploiting this information to split the peptides into groups with similar chromatographic peak widths. Subsequently controlling the FDR separately in each group boosts power relative to an ungrouped approach.

### 2.4 Peptide score normalization

Many search algorithms implement spectrum-level score calibration to normalize scores and boost detection power.^29;33^ This normalization helps boost power because it can account for biases unique to specific spectra (e.g., more sparse spectra might tend to have lower scores). It has been noted that similar biases are present for each peptide, where properties such as length and charge state systematically influence their scores.^34^ Therefore, we aimed to implement peptide-level score normalization to increase detection power. This normalization would not be possible in DDA-type scenarios where sampling for a given peptide will be inherently sparse. However, in the case of DIA data, each peptide’s *m/z* range is systematically sampled across the run, making this type of normalization meaningful. This calibration helps eliminate biases for or against specific subsets of peptides. For example, singly charged peptides tend to have lower scores because they have shorter sequences and therefore yield fewer ions.

Recall that when determining the peak width above we used robust z-score normalization to normalize the rows of the matrix scores *M*. However, the goal here is different: for width-detection the focus of the normalization is on effectively normalizing the relative differences within a row. In contrast, the purpose of peptide normalization is to be able to compare maximal PSM scores across different peptides. Accordingly we found that it is better to use a different, Tailor-like, normalization here. Specifically, analogously to Tailor we normalize scores for a given peptide to the 99th quantile for the peptide. This normalization boosts our power to detect target peptides because the scores are now calibrated between the peptide candidates for each mass spectrum (Supplemental Algorithm 5). Note that this normalization is not optimal for peak-width detection because the additive constant that is added to each score tends to overly-smooth the differences within each row making it difficult to find a uniform cutoff that will define the peak width as effectively as when using robust z-scores.

### 2.5 Selection of optimal representatives

One potential danger in searching DIA data is assigning the same set of fragment ions in a tandem mass spectrum to multiple peptide sequences that happen to share a number of theoretical fragment ions. Although this risk is mitigated by using the narrower 2-*m/z* isolation windows, there are still peptides that fall in the same *m/z* isolation window and share fragment ions. This issue is particularly relevant when searching for post-translationally modified peptides with multiple possible localization sites. In these cases, many possible localization isoforms may score highly when only one is truly present. To address this problem we developed a greedy procedure designed to select the optimal representatives from groups of highly similar (potentially) co-eluting peptides. In the above scenario this means that similar, modified peptides are competing against each other and only the highest scoring modified peptide is retained.

While one can determine the degree of similarity between two peptides by directly comparing their theoretical spectra, this would be a very time consuming job. Therefore, as an alternative we postulate that peptides with shared fragment ions will have highly correlated scores when matched against all tandem mass spectra, or equivalently across all retention times (Supplemental Figure 4). Thus we can assess the similarity between two peptides *p_i_* and *p_j_* by determining the similarity between their corresponding rows of scores *M_i_* and *M_j_*. More specifically, we look at the normalized dot product, or the cosine of the angle, between these two vectors.

In practice our procedure goes through the list of peptides sorted in decreasing order and for each considered peptide *p_i_*, it removes all lower scoring peptides *p_j_* that (i) have overlapping elution profiles, as determined from the chromatographic peak width and boundary selection, (ii) have a statistically significant angle-cosine correlation relative to random peptides, and (iii) share at least 25% of theoretical fragment ions.^35^ This filtering process is applied separately to the target and decoy peptides and is described in detail in Supplemental Algorithm 7.

### 2.6 False discovery rate estimation

Accurate FDR estimation is challenging for DIA data due to the presence of potentially chimeric tandem mass spectra. We aimed to create an FDR control method that will leverage the information provided by DIA data to increase power. Therefore, to control the FDR, we employ a multistep procedure that includes a novel target/decoy peptide competition (TDC) step.

First, we select the maximum score and corresponding spectrum for each peptide. Then, we retain only the highest remaining scoring peptide for each tandem mass spectrum. Although DIA data contains chimeric spectra, it is unlikely that peptides in the same isolation window share the same maximally matched spectrum, especially after representative selection in Section 2.5. In a two hour gradient, we may collect over 150,000 spectra, but detect fewer than 10,000 peptides in most gas-phase fractionated DIA runs. Indeed, we find that using more than one peptide per tandem mass spectrum for FDR control actually reduces power at low false discovery rates possibly due to less-than-perfect score calibration (Supplemental Figure 5).

Each of the peptides remaining after the last step is associated with a score as well as a target/decoy label. While we can pool all of these remaining peptides together to do a standard TDC, grouping peptides according to their chromatographic peak widths and applying TDC separately to each group yielded more discoveries in our analysis. Moreover, when considering PTMs, we can possibly further divide the groups according to the post-translational modification status of the peptides, again followed by FDR control within each group using standard target/decoy procedures described in Supplemental Algorithm 1. Using grouping in this way may allow for making more discoveries,^36;37^ but it can also be detrimental if some groups do not contain a sufficient number of peptides with high scores. Therefore, we advise using caution when a small number of high scoring targets are present in the data. In those cases, the use of groups may reduce discriminatory power by dividing an already low number of high scoring targets into even smaller groups.

## 3 Methods

### 3.1 Data acquisition

Here, we analyze a dataset from Pino *et al*. 2020, ^38^ which is available through the ProteomeXChange Consortium ^39^ with the dataset identifier PXD014815 and Panorama Public^40^ at https://panoramaweb.org/matrix-matched_calcurves.url. A second dataset enriched for phosphorylated peptides was generated for this project. All original data have been deposited to PRIDE Archive (http://www.ebi.ac.uk/pride/archive) with the dataset identifier PXD029733 (Username; Password: lvjPKBfa).

#### 3.1.1 Phosphorylated peptide enriched dataset

The phosphorylated peptide dataset was acquired in house. HEK 293T cells were lysed by resuspending in lysis buffer (8 M Urea, 100 mM Tris pH 8.0, 100 mM NaCl) with Pierce protease and phosphatase inhibitors (Thermo Fisher Scientific). Lysates were incubated on ice for 30 minutes. Lysate protein concentration was measured with a Pierce BCA assay kit (Thermo Fisher Scientific). Lysates were reduced with 5 mM dithiothreitol (DTT) for 30 minutes at 55°C and alkylated with 15 mM iodoacetamide for 30 minutes in the dark, before being quenched with additional 5 mM DTT for 15 minutes.

Lysates were digested on KingFisher Flex robot (Thermo Fisher Scientific) according to R2-P1 protocol described in Leutert et al., 2019.^41^ In brief, magnetic beads (MagReSyn Amine, ReSyn Biosciences) were combined with lysates in a 96-well plate at a 10:1 w/w ratio of bead to protein. Ethanol was added to bead/lysate mixture to a final concentration of 70%. Samples were then washed by transferring beads to a wash plate containing 80% ethanol. This wash was repeated two more times. Proteins were digested by transferring beads to plate containing trypsin (1:50) and ammonium bicarbonate pH 8.2. Samples were incubated for 6 hours at 37° C. Beads were then removed and the remaining solvent was evaporated using vacuum centrifugation. Samples were enriched for phosphorylated peptides as described in Leutert et al.^41^ In brief, peptides were enriched on KingFisher Flex using 25 *μL* of 5% Fe^3+^NTA magnetic beads (Cube Biotech) per 100 *μg* of peptide. Samples were reconstituted in 80% acetonitrile (ACN), 0.1% trifluoroacetic acid (TFA) and incubated with beads for 30 minutes before being washed three times with 50% ACN, 0.1% TFA. Phosphorylated peptides were then eluted from beads with 50% ACN, 2.5% NH_4_OH. Phosphorylated peptides were then dried in a speed vacuum resuspended in 3% ACN 5% formic acid for LC-MS/MS analysis.

Data were acquired using on an Orbitrap Exploris 480 (Thermo Fisher Scientific) coupled to an Easy-nLC 1200 (Thermo Fisher Scientific). Peptides were separated on an in-house pulled 100 *μm* inner diameter C18 column packed with 30 cm of 3 *μm* beads (Dr. Maisch GmbH) using a 90-min gradient from 3 to 32% ACN with 0.1% formic acid. Mass spectrometry was performed as DIA gas-phase fractions with six fractions. Each fraction covered a full MS scan range of 100 *m/z* from 400-1000 in total (400-500, 500-600, 600-700, 700-800, 800-900, 900-1000). Full MS scans were acquired at 30,000 resolution with an AGC target of 100% and a maximum injection time of 100 ms. MS/MS spectra were acquired with staggered 4 *m/z* windows at 30,000 resolution, AGC target of 1000%, maximum inject time of 25 ms, and a HCD collision energy of 27%.

#### 3.1.2 Yeast dataset

The *S. cerevisiae* dataset comes from Pino et al. 2020.^38^ From this dataset, a chromatogram library generated from data collected in 6 gas-phase fractions was used along with quantitative DIA data on three 100% and three 50% yeast samples. Yeast strains BY4741 and S288C were cultured in both YEPD and ^15^N minimal media and harvested at mid-log phase. The cells were lysed with bead beating in 8 M urea buffer and the lysates were reduced, alkylated, and digested with trypsin for 16 hours. The peptides were desalted with a mixed-mode (MCX) method and synthetic retention time standards were added. Dilutions were achieved by diluting cultures from the YEPD media in cultures grown in the ^15^N minimal media.

For LC-MS/MS analysis, peptides were separated via reversed-phase liquid chromatography using a Waters NanoAcquity ultra-performance liquid chromatography (UPLC) with a 30 cm long 75 *μm* inner diameter fused-silica capillary column packed with 3 *μm* ReproSil-Pur C18 beads (Dr. Maisch GmbH, Ammerbuch, Germany). Trapping was performed with a 150 *μm* fritted capillary. Solvent A was 0.1% formic acid in water and solvent B was 0.1% formic acid in 98% acetonitrile. Each injection consisted of a 90-minute gradient from 5 to 35% B, followed by a 10-minute ramp to 60% B, a wash step (5-minute ramp to 95% B and a 5-minute wash at 95% B), and a re-equilibration step (ramp down to 2% B for 1 minute and hold at 2% B for 19 minutes).

Tandem mass spectrometry data were acquired using DIA performed on a Thermo Q-Exactive HF Orbitrap mass spectrometer monitoring a mass range of 388.43190–1012.70480 *m/z* using normalized collision energy of 27 with assumed charge state of +2. Gas-phase fractionated “narrow-window” DIA data were acquired on a sample of yeast grown in YEPD media with the following parameters: 4 *m/z* staggered DIA windows, 30,000 resolution, 55 ms maximum inject time, and 1e6 AGC target. For the quantitative data, 24 *m/z* overlapping DIA windows were acquired using the same resolution, maximum inject time, and AGC as the narrow-window DIA data.

### 3.2 Data processing

All raw files are preprocessed to demultiplex overlapping windows using ProteoWizard’s MSConvert (v 3.0.1908) using vendor peak picking, overlap-only demultiplexing with a mass error of 10 ppm, and the “SIM as spectra” option turned on.^42^ All spectra are assigned possible charge states +1, +2, and +3. Database searches are performed in Crux Tide (v 3.2)^27^ using a precursor tolerance of 1.007 m/z and Tailor calibration.^29^ All searches include carbadomethylation of cysteine as a fixed modification and appropriate post-translational modifications or missed cleavages. All processing and analysis were performed in Python (v 3.8.2) and R (v 3.6.1). Scripts used for processing and analysis are available as supporting information.

#### 3.2.1 Analysis of phosphorylated peptide-enriched dataset

For the phosphorylated peptide search, the human protein FASTA database was downloaded from Uniprot (UP000005640, https://www.uniprot.org/) in July of 2019. Prior to database searching, a Tide index was constructed by performing an *in silico* digestion of the FASTA with trypsin. The index contained fully tryptic peptides with no missed cleavages and exactly one variable modification of 79.966331 on serine, threonine, or tyrosine (Supplemental Table 2). Using this index, a Tide search of the phosphorylated peptide-enriched DIA data was performed in Crux as described in the previous section. Specific search parameters are listed in Supplemental Table 3. The results from this Tide search were processed according to the procedure described in Section 2. No groups were used for FDR estimation due to the small number of high scoring peptides in possible groups.

For DIA-Umpire analysis, pseudo-spectrum extraction was performed on demultiplexed mzXML files with DIA-Umpire (V2.0)^8^ using window-type MSX. A full list of these extraction parameters are listed in Supplemental Table 4. A Tide search was performed on the resulting pseudo-spectra using the same index as the direct search (Supplemental Table 2) with a precursor tolerance of 20 parts per million and a top-match of three (Supplemental Table 5). FDR estimation was performed using Crux Percolator^43^ with all default settings. Peptide-level q-values from this search were used for further analysis.

#### 3.2.2 Analysis of *S. cerevisiae* dataset

The FASTA database for *S. cerevisiae* strain ATCC 204508 was downloaded off of Uniprot (UP000002311, https://www.uniprot.org/) in September of 2020. A Tide index was constructed from the *S. cerevisiae* proteome using default settings for tryptic peptides (Supplemental Table 6). A Tide search was performed on the gas-phase fractionated yeast dataset^38^ with a top match of 10,000 (Supplemental Table 7). All peptides with q-values below 1% were converted to SSL format and built into a spectral library using Bibliospec^44^ in Skyline. The spectral library .blib file was converted to an EncyclopeDIA-compatible format in EncyclopeDIA for later use to search quantitative DIA data.

The predicted spectral library was made by generating a list of all fully tryptic unmodified peptides from *S. cerevisiae* in EncyclopeDIA and predicting their spectra using the Prosit pre-trained spectral prediction model accessed through the online server (https://www.proteomicsdb.org/prosit/).21A chromatogram library was generated from the narrow window DIA data using EncyclopeDIA (v 0.9.0)^13^ with a normal target/decoy approach requiring at least three quantitative ions. All six gas-phase fractionated DIA runs were searched individually and combined in EncyclopeDIA to generate a chromatogram library .elib file.

Both the spectral library from our workflow and chromatogram library from the Prosit search were used to search quantitative wide-window DIA data in EncyclopeDIA^13^. All six quantitative DIA mass spectrometry runs were searched against each library and the results were combined using the “Save Quant Reports” option in EncyclopeDIA. Peptides accepted at 1% EncyclopeDIA FDR with at least five quantitative transitions were used for quantitative analysis. Area under the curve data calculated by EncyclopeDIA was analyzed in R to assess fold-change abundances.

## 4 Results and discussion

### 4.1 The proposed workflow increases sensitivity for detecting phosphorylated peptides relative to pseudo-spectrum approach

To assess the ability of our workflow to detect post-translationally modified peptides by generating libraries in cases where spectral libraries are not readily available, we sought to detect phosphorylated peptides in DIA data from HEK 293T cells. While there are several methods for the detection of phosphorylated peptides in DIA data, most rely on the use of existing libraries.^45^Although there are high-quality spectral libraries available for phosphorylated peptides,^46–48^that is not the case for many other post-translational modifications.

As a reference point, we used DIA-Umpire^8^ as an alternative library-free approach. Specifically, pseudospectra were extracted using DIA-Umpire^8^ and searched using Tide (with Percolator used only for false discovery rate estimation). These results provide a proof-of-concept that our workflow offers a powerful alternative method for detecting PTMs directly from DIA mass spectrometry data.

Overall, our direct search workflow detects more peptides at every confidence level relative to the DIA-Umpire approach (Figure 2A). At an FDR threshold of 1%, our workflow detects 1300 unique peptides while the DIA-Umpire approach detects 851 peptides. We observe overlap between the phosphorylated peptides detected by both searches at a 1% FDR (Figure 2B) although our workflow is able to detect many more unique peptides. These results suggest that our workflow is a viable alternative to other library-free methods for detecting PTMs in DIA data.

**Figure 2:**
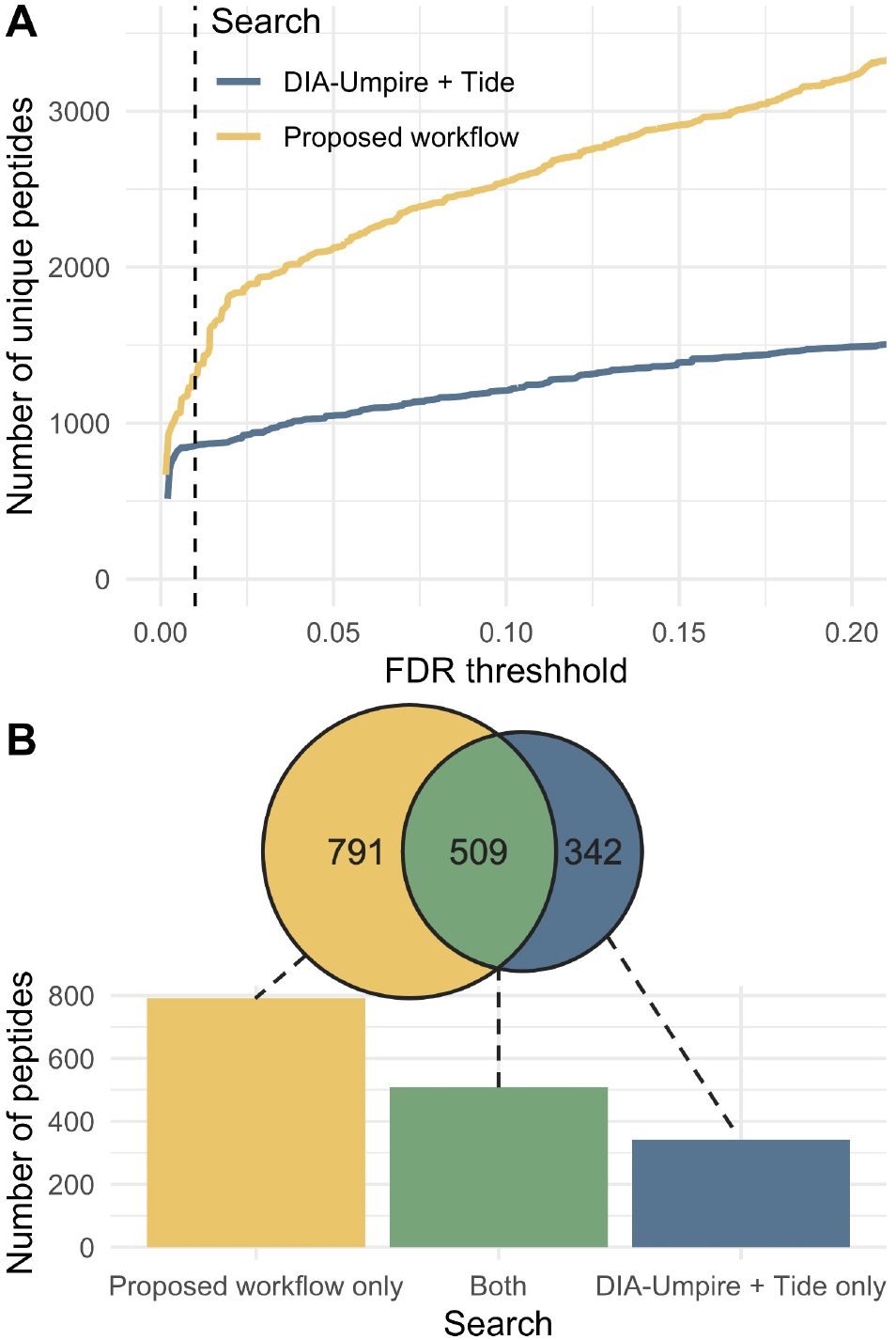
Comparison of searches for phosphorylated peptides. (A) Plot of number of peptides detected at a given FDR threshhold obtained by searching for doubly charged phosphorylated peptides using our workflow compared to a search with DIA-Umpire results with Tide and adjusting FDR with Percolator. (B) The 1% FDR threshold is indicated with a dashed vertical line. The number of peptides unique to each search as well as those shared by both methods are also displayed at a 1% FDR threshold.

One reason our workflow is more sensitive in detecting phosphorylated peptides than the pseudo-spectrum based approach (Figure 2A) is that it is independent of signal in the precursor (MS1) spectra. Due to automatic gain control in trapping-type instruments, MS1 signal can be less sensitive than MS2.^10;11^ DIA-Umpire and similar methods are optimized to detect peptides in wide-window DIA data where the MS2 signal may be less sensitive than it is with narrow isolation windows due to limits on total ion signal. Therefore, our workflow is more sensitive due to its ability to detect target peptides without an MS1 signal, while DIA-Umpire cannot. As further evidence of the differences in these methods, both methods detect a large number of unique peptides at 1% FDR (Figure 2B). This finding suggests that the two methods use orthogonal methods to detect peptides. While the DIA-Umpire approach may be more sensitive in some cases, the fact that our workflow does not rely on MS1 signal helps it perform better for narrow-window DIA mass spectrometry data.

### 4.2 Peptides detected with the proposed workflow are quantitatively accurate

To ensure that our workflow is producing accurate results, we assessed the quantitative accuracy of peptides detected with our workflow compared to EncyclopeDIA, a peptide-centric method for peptide detection in DIA data.^13^ We created a library of all peptides detected with our workflow at 1% FDR in narrow-window DIA data. We compared these results to a library generated by searching the narrow-window data in EncylopeDIA^13^ using a Prosit predicted library.^21^We then used both libraries to search quantitative wide-window DIA data of 100% and 50% yeast in YEPD media diluted with yeast grown in ^15^N media and compared quantitative accuracy of all peptides.

In this analysis, we use the ratio of signal between the two datasets as “ground truth” to evaluate search accuracy. If a peptide is really present, we would expect to see a two-fold decrease in signal in the 50% sample relative to the 100% sample, which corresponds to a quantitative ratio of two. Peptides detected in both searches were highly accurate (Figure 3B) with a median quantitative ratio of 2.14. Peptides unique to the library generated with the proposed workflow and the predicted library had median quantitative ratios of 2.02 and 2.31, respectively.

**Figure 3:**
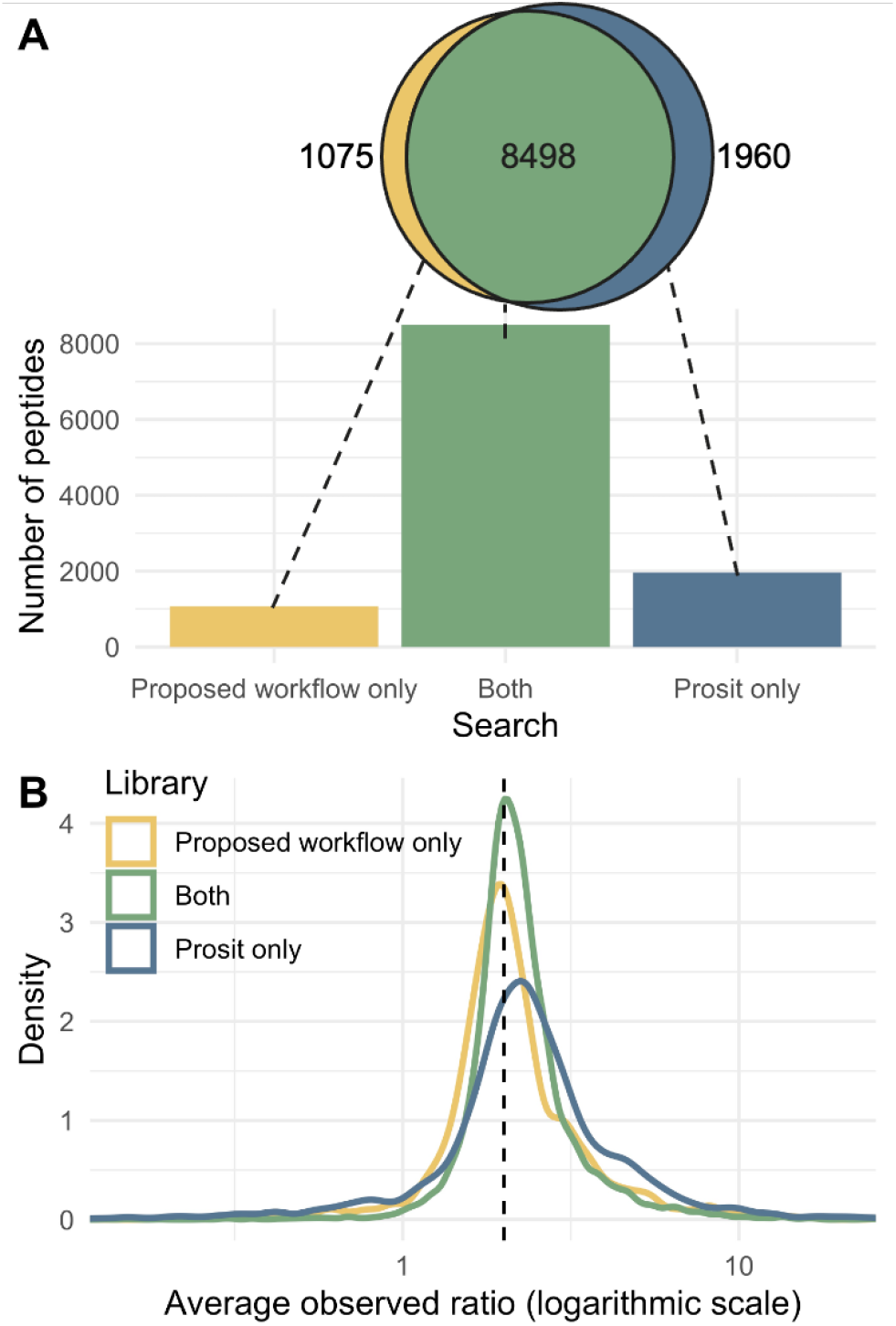
Comparison of peptides detected in quantitative data. Quantitative peptides were detected in the EncyclopeDIA search of wide-window DIA data using both a library generated from our workflow and a narrow-window calibrated Prosit predicted library. The overlap between peptides detected with both methods at a 1% FDR threshold (A) is shown along with a normalized density plot of the observed ratio of the peptides in each group between an undiluted sample and a sample diluted 50% (B). For the quantitative comparison, the expected ratio of 2 is shown as a dashed vertical line.

The library created by our direct search detects similar, but slightly lower, numbers of quantitative peptides relative to a Prosit predicted library in the *S. cerevisiae* dataset (Figure 3). At 1% FDR, both searches have a high degree of overlap in quantitative peptides as determined by EncyclopeDIA (Figure 3A). These two searches have much higher overlap than the previous analysis (Figure 2B) because neither rely on MS1 signal. Although the Prosit library is able to detect 9% more quantitative peptides, the peptides unique to that search are less accurate than the peptides found in both searches and those unique to the direct search (Figure 3B). Therefore, the extra detections in the Prosit search may in part be due to a higher false-positive rate. These results suggest that our search workflow compares favorably to using another popular library-free DIA search method, using EncyclopeDIA as a peptide-centric DIA approach to search a predicted spectral library.

We used a 1% FDR threshold to generate our library here, but one could also use a higher FDR threshold (Supplemental Figure 6). However, the higher cutoff comes with an increased risk of false detections, and we therefore suggest using the conservative 1% threshold. We note that the 1% FDR cutoff library is highly efficient in the sense that we are detecting a much larger fraction in the EncyclopeDIA quantitative search than the narrow-window calibrated Prosit library (Supplemental Figure 6). At the same time this gives further evidence of the accuracy of our search.

## 5 Conclusions

Here, we propose a library-free workflow to detect peptides in narrow-window DIA data without relying on MS1 signal. We address many of the issues in searching for PTMs in DIA data, including the reliance on a spectral library. Existing spectral libraries may not contain all relevant peptides and may not be available for all PTMs. Additionally, spectral and retention time prediction methods are not trained for many PTMs. Our method is able to detect PTMs from narrow-window DIA mass spectrometry data to create libraries that may be used for quantitative DIA mass spectrometry analysis.

The gas-phase fractionated DIA data used here could easily be collected in almost any experiment with minimal sample use and instrument time compared to deep-fractionation DDA-based libraries. These results show that this type of search is a promising avenue for future development, but it will require computational adjustments to become readily useable for many applications. While DIA has many advantages over DDA methods, its uses can be limited by computational hurdles. Researchers wishing to study non-model organisms and/or specific post-translational modifications may be unable to rely on many existing tools and spectral libraries. Therefore, this type of search workflow is crucial for moving the field forward and making DIA accessible to a broader range of applications.

## 6 Acknowledgments

This work is supported in part by National Institutes of Health Grants P41 GM103533, U01 DK121289, and U19 AG065156.

## 7 Notes

The authors declare the following competing financial interest(s): The MacCoss Lab at the University of Washington has a sponsored research agreement with Thermo Fisher Scientific, the manufacturer of the instrumentation used in this research. M.J.M. is a paid consultant for Thermo Fisher Scientific.

## 8 Supporting Information

- **Supplemental File S1:** PDF containing pseudocode for the search procedure along with supplemental tables and figures.
- **Supplemental File S2:** Python script (precursor_matrix.py) used to convert Tide search results to matrix format.
- **Supplemental File S3:** R script (process_precursors.R) used to perform search.
- **Supplemental File S4:** Python script (precursor_confidence.py) used for FDR control.

---

For TOC Only

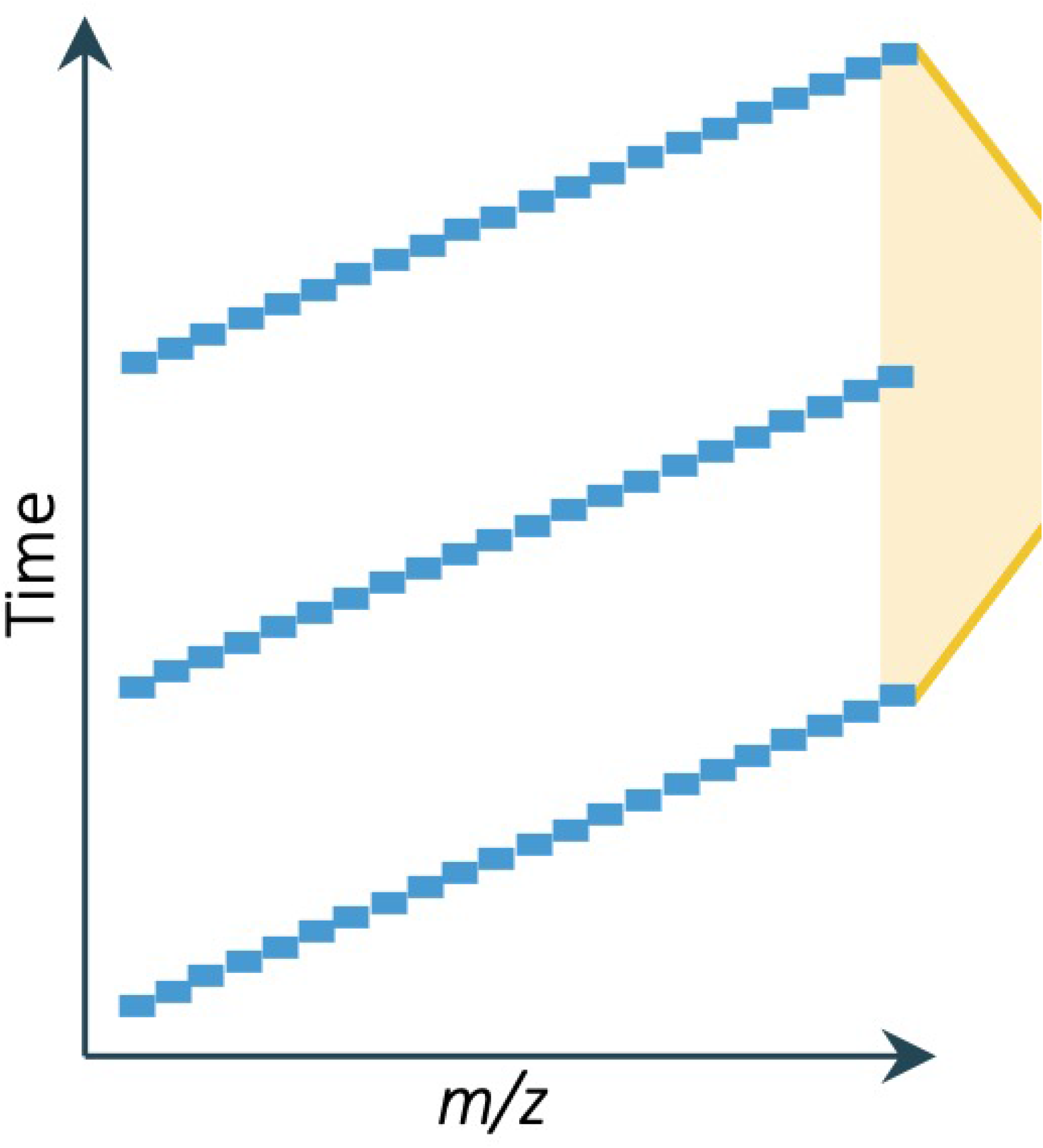

