## Supplemental File S1 for "A flexible workflow for building spectral libraries from narrow window data independent acquisition mass spectrometry data"

### 1 Supporting Information

- **Supplemental File S1:** PDF containing pseudocode for the search procedure along with supplemental tables and figures.
- **Supplemental File S2:** Python script (precursor_matrix.py) used to convert Tide search results to matrix format.
- **Supplemental File S3:** R script (process_precursors.R) used to perform search.
- **Supplemental File S4:** Python script (precursor_confidence.py) used for FDR control.

| | |
|---|---|
| $allS$ | set of MS2 spectra structured as a list of arrays, one for each isolation window |
| $allD$ | database of all possible peptides (where each peptide is a specific peptide/charge state combination) structured as a list of arrays, each corresponding to one isolation window |
| $D$ | database of all possible peptides within a single precursor isolation window |
| $S$ | all MS2 spectra associated with a single precursor isolation window |
| $P$ | all possible peptides associated with a single isolation window |
| $m$ | number of peptides |
| $n$ | number of MS2 spectra |
| $p_i$ | a single peptide |
| $s_j$ | a single MS2 spectrum |
| $M_{i,j}$ | score assigned to the match between peptide $p_i$ and spectrum $s_j$ |
| $A$ | vector of changepoints |
| $peak$ | an m-dimensional structure array which stores the following information for each peptide: the maximally matching spectrum ($peak.m$), the corresponding score ($peak.s$), the width of the peak ($peak.w$), and its left and right boundaries ($peak.l$, $peak.r$) |
| $TDpairs$ | a data structure containing target/decoy pairing information |
| $groups$ | desired grouping designations for FDR control |
| $isTarget$ | a logical vector indicating the target/decoy status of each peptide in the isolation window |
| $Ip$ | indices of considered peptides |
| $\tau$ | acceptance threshold |
| $allGroups$ | desired grouping aggregated across all precursor isolation windows |
| $allIsTarget$ | target/decoy identification aggregated across all precursor isolation windows |
| $allScores$ | assigned peptide scores aggregated across all precursor isolation windows |
| $allPs$ | IDs of all considered peptides aggregated across all precursor isolation windows |
| $qvalue$ | q-values of all considered target peptides aggregated across all precursor isolation windows |

**Table 1: Notation**

**Algorithm 1 DIA analysis**

```
procedure diaSearch(allS, allD, groups)
    allGroups[1 : length(groups)] = [ ]
    allIsTarget = [ ]
    allScores = [ ]
    allPs = [ ]
    for i_w = 1 : length(allS) do                                  ▷ loop on isolation windows
        D_T := allD[i_w]; S := allS[i_w]
        [M, TDpairs, isTarget] := databaseSearching(D_T, S)
        D_ext = [D_T, D_T]            ▷ D_ext includes precursor ID of targets and corresponding decoys
        M := changepointDetection(M)        ▷ Remove leading and trailing “junk” spectra from M
        peak := chromatographicPeakDetection(M, TDpairs, isTarget)
        M := peptideScoreNormalization(M)
        Ip := TDC(M, TDpairs, peak, isTarget)       ▷ keep the higher scoring peptide from each target-decoy pair
        Ipt := optimalRepresentativeSelection(M, Ip, isTarget, peak)
        Ipd := optimalRepresentativeSelection(M, Ip, NOT(isTarget), peak)
        Ip := [Ipt, Ipd]
        mAll := 0                          ▷ mAll is the windows-aggregated number of peptides
        for i_p in Ip do
            if i_p ≠ which.max(M[:, peak[i_p].m]) then
                Ip = Ip[−i_p]                    ▷ Retain only the maximum peptide per spectrum
            end if
            mAll := mAll + 1
            i_g = whichGroup(D_ext[i_p], isTarget[i_p], peak[i_p], groups)     ▷ Determine the group of i_p based on its features
            allGroups[i_g] = [allGroups[i_g], mAll]
        end for
        allPs = [allPs, D_ext[Ip]]
        allScores = [allScores, peak[Ip].s]
        allIsTarget = [allIsTarget, isTarget[Ip]]
    end for
    for i_g = 1 : length(groups) do
        Ip = allGroups[i_g]
        qvalues[Ip] = qvaluesViaTDC(allScores[Ip], allIsTarget[Ip])
    end for
    return (allPs[allIsTarget], qvalue[allIsTarget], allScores[allIsTarget])
end procedure
procedure qvaluesViaTDC(scores, isTarget)
    n := length(scores)
    sortPerm := order(scores)
    scores := scores[sortPerm]                               ▷ sort scores in decreasing order
    isTarget := isTarget[sortPerm]
    nTargetWins := cumsum(isTarget)
    nDecoyWins := [1 : n] − nTargetWins
    estFDR := min (1, (nDecoyWins + 1)/ max (1, nTargetWins))
    qvalues[n] := estFDR[n]
    for i = n − 1 : 1 by −1 do
        qvalues[i] := min (estFDR[i], qvalues[i + 1])
    end for
    return (qvalues[inversePermutation(sortPerm)])
end procedure
```

**Algorithm 2** Database searching

**procedure** DATABASESEARCHING($D_T = (p_i)_1^m,\ S = (s_j)_1^n$)
  $D_D$ = CREATEDECOYDB($D_T$)
  **for** $1 \le j \le n$ **do**
    $M_T[i, :] :=$ SCOREALLPEPTIDES($s_j$, $D_T$) ▷ SCOREALLPEPTIDES returns the scores of the matches between the spectrum $s_j$ and every peptide $p_i \in D$ (here we used Tailor-normalized XCorr)
    $M_D[i, :] :=$ SCOREALLPEPTIDES($s_j$, $D_D$)
  **end for**
  $isTarget[1 : m]$ = TRUE
  $isTarget[m + 1 : 2m]$ = FALSE
  $TDpairs[1 : m] = [m + 1 : 2m]$
  $TDpairs[m + 1 : 2m] = [1 : m]$
  **return** [$M :=$ CONCAT($M_T$, $M_D$), $TDpairs$, $isTarget$]
**end procedure**

**Algorithm 3** Changepoint detection

**procedure** CHANGEPOINTDETECTION($M$)
  $n$ = NROWS($M$)
  **for** $j = 1 : n$ **do**
    $meds[j] :=$ MEDIAN($M[:, j]$)
  **end for**
  **for** $ncp = 2 : 4$ **do** ▷ $ncp$ is total number of changepoints
    $A :=$ CHANGEPOINT($meds$, $ncp$) ▷ Inputs for changepoint function (described by Killick et al., 2016) are vector of scores and number of changepoints, $l$
    **if** $A_{ncp} - A_1 \ge 0.5 \times n$ **then**
      $M := M[:, A_1 : A_{ncp}]$
      **return** $M$
    **end if**
  **end for**
  **return** `ERROR`
**end procedure**

**Algorithm 4 Chromatographic peak detection**

```
procedure chromatographicPeakDetection(M, TDpairs, isTarget)
    m = nrows(M)
    for i = 1 : m do
        med = median(M[i, :])
        MAD = median(|M[i, :] − med|)
        M^RZ[i, :] := (M[i, :] − med)/MAD
    end for
    for i_t = 1 : m do
        if isTarget[i_t] then
            i_d = TDpairs[i_t]
        else
            continue
        end if
        [peak[i_t].m, l_t, r_t] := findPeakInRow(M^RZ[i_t, :])
        [peak[i_d].m, l_d, r_d] := findPeakInRow(M^RZ[i_d, :])
        if l_t + r_t < l_d + r_d then
            l = l_d; r = r_d
        else
            l = l_t; r = r_t
        end if
        peak[i_t].l = max (peak[i_t].m − l, 1)
        peak[i_t].r = min (peak[i_t].m + r, n)
        peak[i_t].w = l + r + 1
        peak[i_t].s = M[i_t, peak[i_t].m]
        peak[i_d].l = max (peak[i_d].m − l, 1)
        peak[i_d].r = min (peak[i_d].m + r, n)
        peak[i_d].w = l + r + 1
        peak[i_d].s = M[i_d, peak[i_d].m]
    end for
    return peak        ▷ an m-dimensional structure array storing peak data for each peptide
end procedure
procedure findPeakInRow(M_r)
    m = which.max(M_r)
    M_r = M_r/M_r[m]
    for l = 0 : m − 1 do
        if M_r[m − l] < 0.75 then
            l = l − 1
            break
        end if
    end for
    for r = 0 : n − m do
        if M_r[m + r] < 0.75 then
            r = r − 1
            break
        end if
    end for
    return [m, l, r]
end procedure
```

**Algorithm 5 Peptide score normalization**

```
procedure peptideScoreNormalization(M)
    m = nrows(M)
    for i = 1 : m do
        q^0.99 := Quantile(M[i, :], 0.99)
        M[i, :] = M[i, :] / q^0.99
    end for
    return M                                ▷ return the peptide normalized scores
end procedure
```

**Algorithm 6 Target/decoy competition**

```
procedure TDC(M, TDpairs, peak, isTarget)
    m = nrows(M)
    Ip := ∅
    for i_t = 1 : m do
        if isTarget[i_t] then
            i_d = TDpairs[i_t]
            if peak.s[i_t] > peak.s[i_d] then
                Ip = [Ip, i_t]
            else
                Ip = [Ip, i_d]
            end if
        end if
    end for
    return Ip
end procedure
```

**Algorithm 7 Selection of optimal representatives**

**procedure** OPTIMALREPRESENTATIVESELECTION($M$, $Ip$, $isTarget$, $peak$)
    $Ipt := Ip[isTarget]$
    $Ipd := Ip[\text{NOT}(isTarget)]$
    $Ipt := IIpt[\text{ORDER}(peak.s[Ipt])]$ ▷ Sort scores in decreasing order
    **for** $i_p$ in $Ipt$ **do**
        $pRow := M[i_p, :]$
        $overlaps = \{i \in Ipt : peak[i_p] \cap peak[i] \neq \emptyset \text{ AND } peak.s[i] < peak.s[i_p]\}$
        $acosNull = 0$
        $draws := \text{PERMUTE}(Ipd)$
        **for** $l = 1 : \min(1000, \text{LENGTH}(Ipd))$ **do**
            $pRow2 = M[draws[l], :]$
            $acosNull := \max(acosNull, (|pRow \cdot pRow2|)/(\|pRow\|\|pRow2\|))$
        **end for**
        **for** $i$ in $overlaps$ **do**
            $pRow2 = M[i, :]$
            $acosTest = (|pRow \cdot pRow2|)/(\|pRow\|\|pRow2\|)$
            **if** $acosTest > acosNull$ AND $[(pRow2 > 0) \cdot (pRow > 0)/n] > 0.25$ **then** ▷ Check if acos distribution and predicted ion overlap are significant
                $Ipt = Ipt[-which(Ipt == i)]$ ▷ Remove the lower scoring peptide from the final matrix
            **end if**
        **end for**
    **end for**
    **return** $Ipt$
**end procedure**

| Parameter | Value |
|---|---|
| min-length | 6 |
| max-length | 50 |
| min-mass | 200 |
| max-mass | 7200 |
| enzyme | trypsin |
| deisotope | 0 |
| digestion | full-digest |
| missed-cleavages | 0 |
| keep-terminal-aminos | NC |
| num-decoys-per-target | 1 |
| min-mods | 1 |
| max-mods | 1 |
| mods-spec | 1STY+79.966331 |

**Table 2: Parameters for Tide index of phosphopeptide enriched samples.**

| Parameter | Value |
|---|---|
| min-peaks | 20 |
| deisotope | 0 |
| precursor-window | 1.007 |
| precursor-window-type | mz |
| mz-bin-width | 0.02 |
| mz-bin-offset | 0.4 |
| spectrum-charge | 2 |
| top-match | 100000 |
| use-tailor-calibration | true |
| concat | true |

**Table 3: Parameters for direct Tide search of phosphopeptide enriched samples.**

| Parameter | Value |
|---|---|
| RPmax | 25 |
| RFmax | 300 |
| CorrThreshold | 0.2 |
| DeltaApex | 0.6 |
| RTOverlap | 0.3 |
| AdjustFragIntensity | true |
| BoostComplementaryIon | true |
| ExportPrecursorPeak | false |
| ExportFragmentPeak | false |
| SE.MS1PPM | 20 |
| SE.MS2PPM | 40 |
| SE.SN | 2 |
| SE.MS2SN | 2 |
| SE.MinMSIntensity | 5 |
| SE.MinMSMSIntensity | 1 |
| SE.MaxCurveRTRange | 1 |
| SE.Resolution | 15000 |
| SE.StartCharge | 2 |
| SE.EndCharge | 3 |
| SE.MS2StartCharge | 2 |
| SE.MS2EndCharge | 3 |
| SE.NoMissedScan | 1 |
| SE.MinFrag | 10 |
| SE.EstimateBG | true |
| SE.MinNoPeakCluster | 1 |
| SE.MaxNoPeakCluster | 3 |
| SE.StartRT | 0 |
| SE.EndRT | 9999 |
| SE.MinMZ | 200 |
| SE.IsoPattern | 0.8 |
| SE.MassDefectFilter | true |
| WindowType | MSX |

**Table 4: DIA-Umpire parameters for generating pseudospectra from phosphopeptide-enriched data.**

| Parameter | Value |
|---|---|
| min-peaks | 20 |
| deisotope | 0 |
| precursor-window | 20 |
| precursor-window-type | ppm |
| mz-bin-width | 0.02 |
| mz-bin-offset | 0.4 |
| spectrum-charge | 2 |
| top-match | 3 |
| use-tailor-calibration | true |
| concat | true |

**Table 5: Parameters for Tide search of pseudospectra from phosphopeptide enriched samples.**

| **Parameter** | **Value** |
|---|---|
| min-length | 6 |
| max-length | 50 |
| min-mass | 200 |
| max-mass | 7200 |
| enzyme | trypsin |
| deisotope | 0 |
| digestion | full-digest |
| missed-cleavages | 0 |
| keep-terminal-aminos | NC |
| num-decoys-per-target | 1 |
| min-mods | 0 |
| max-mods | 255 |

**Table 6: Parameters for Tide index for yeast search.**

| **Parameter** | **Value** |
|---|---|
| min-peaks | 20 |
| deisotope | 0 |
| precursor-window | 1.007 |
| precursor-window-type | mz |
| mz-bin-width | 0.02 |
| mz-bin-offset | 0.4 |
| spectrum-charge | all |
| top-match | 10000 |
| use-tailor-calibration | true |
| concat | true |

**Table 7: Parameters for direct Tide search of yeast samples.**

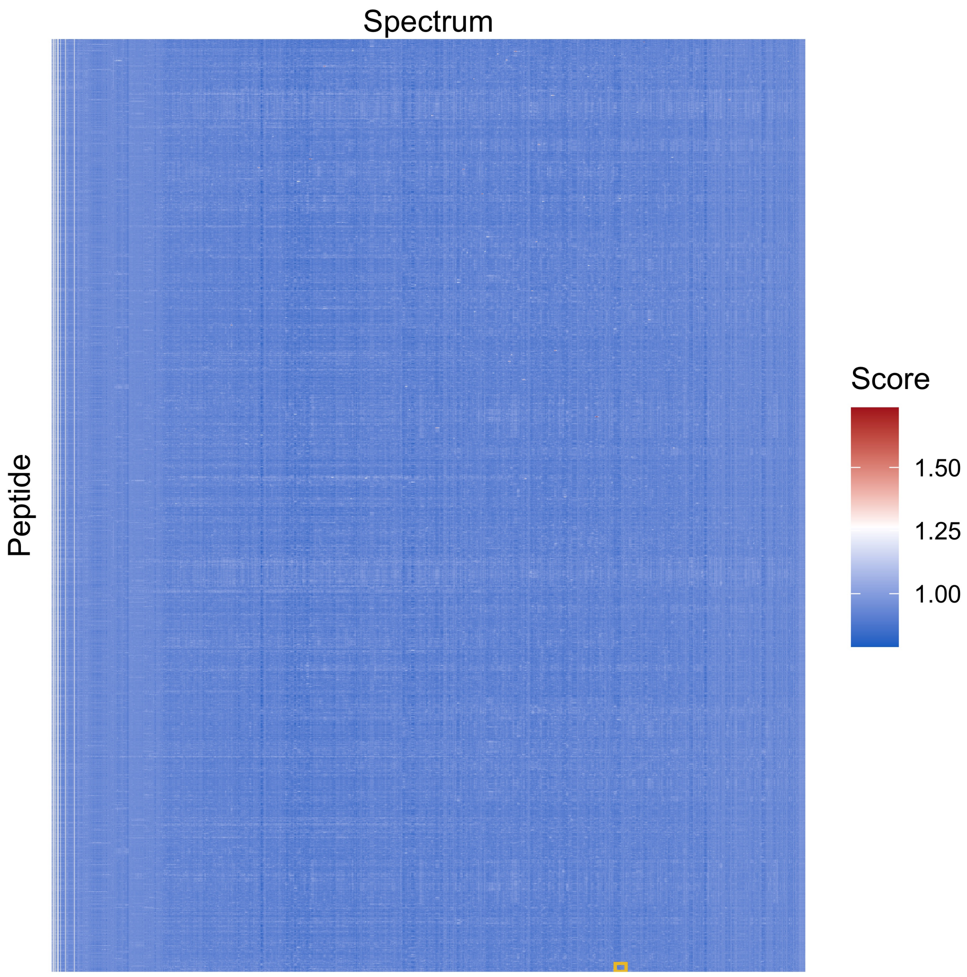


**(a)** Full matrix

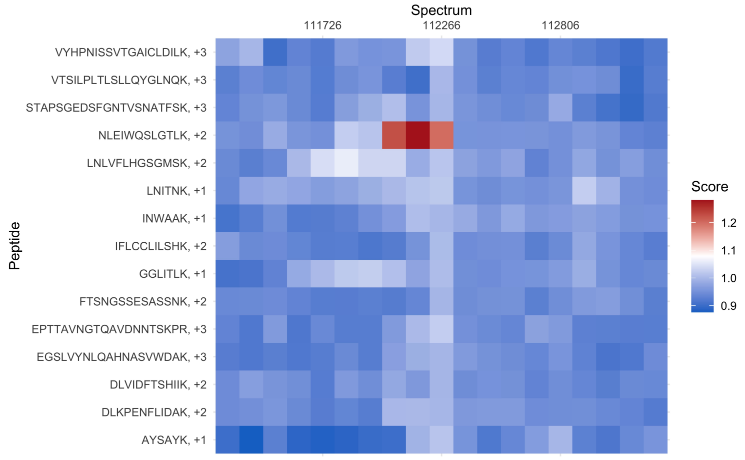


**(b)** Zoom in of (a)

**Figure 1: Graphical representation of a single 2-$m/z$ matrix produced after Tide search** (A) The full matrix has 1733 peptides (rows) and 1439 spectra (columnns). (B) A zoom in on the area of (A) marked by the yellow box. The peptide NLEIQQSLGTLK is accepted at 1% FDR based on this example.

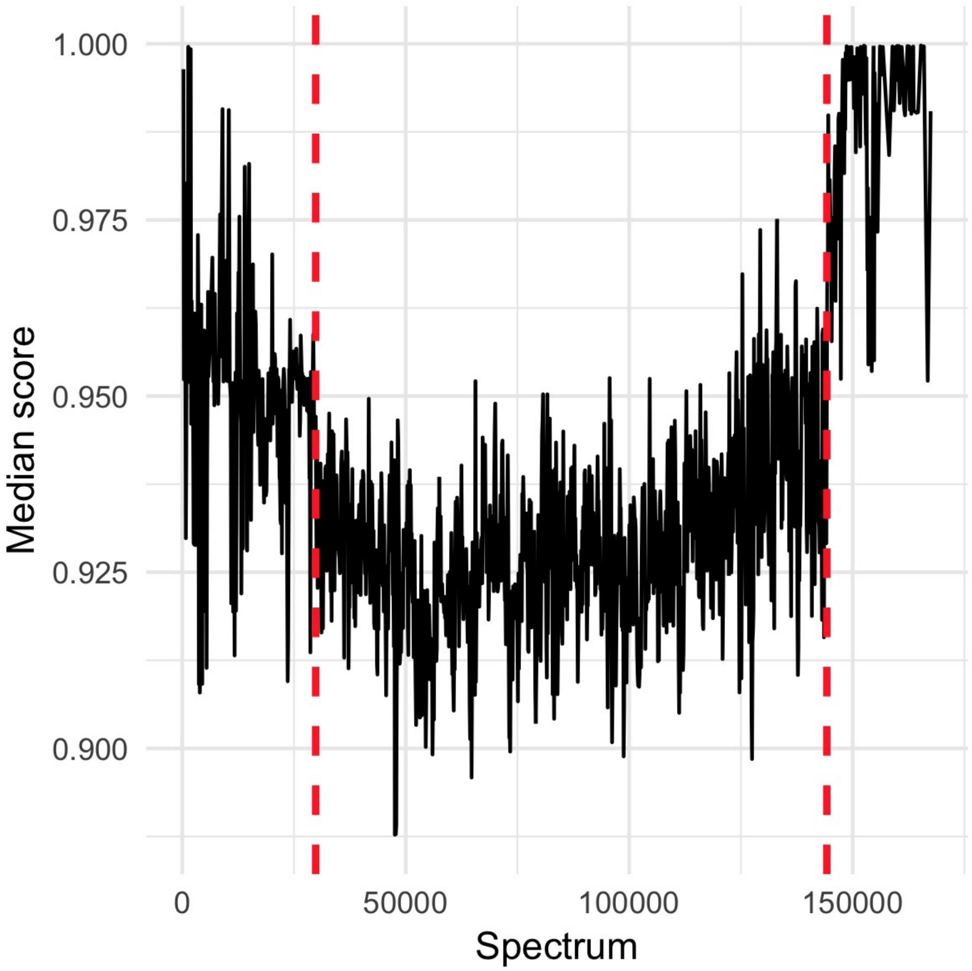


**Figure 2: Plot of median scores for each spectrum across a DIA window.** Changepoint detection automatically removes spectra with median scores distinct from the rest of the run. To account for shifts in score distributions early in the chromatographic gradient, only spectra between red dashed lines are retained for analysis. We first identify two changepoints and retain only the $n'$ mass spectra between these two changepoints provided $n' \geq 0.5n$, where $n$ is the original number of spectra. If $n' < 0.5n$ then we repeat the changepoint detection while increasing the number of changepoints, this time considering the $n'$ spectra between the first and the last changepoints.

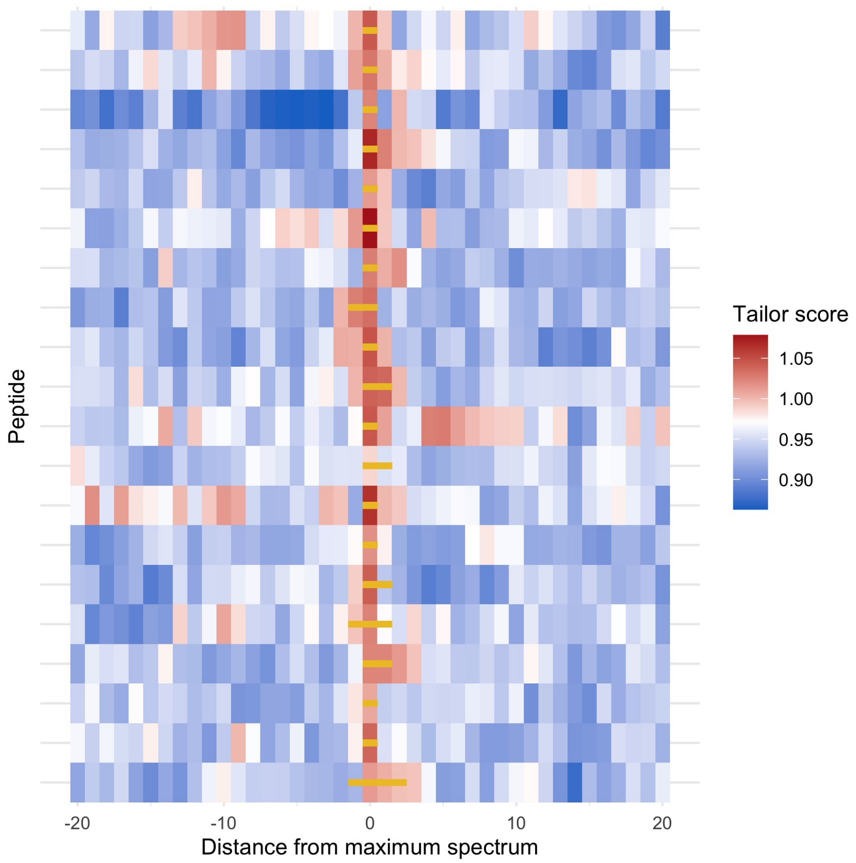


**Figure 3: Peak boundaries assigned with our method.** The figure graphically depicts the +/- 20 scores surrounding the top scoring spectrum for a number of target peptides. Peak boundaries (assigned by Supplemental Algorithm 4) are shown with yellow lines.

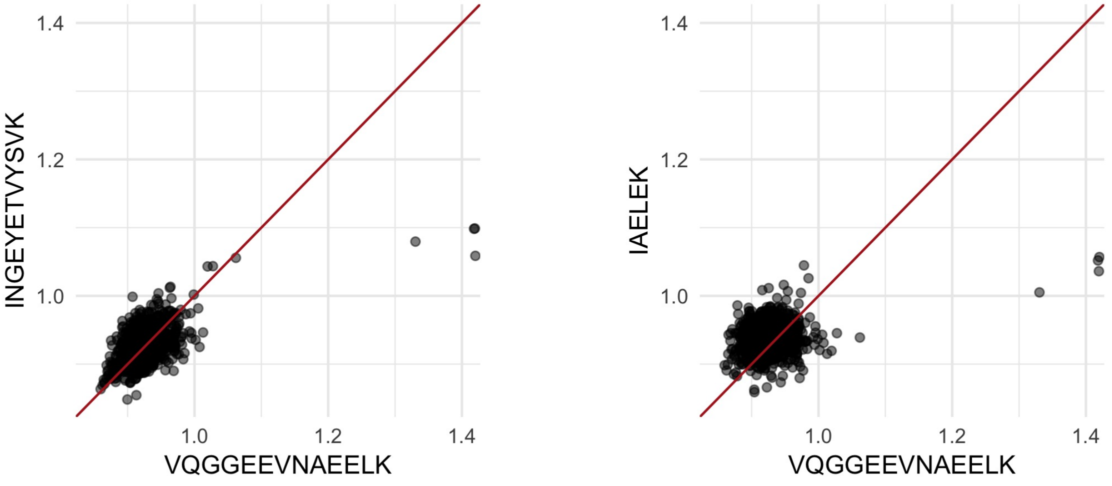


**(a)** Significantly correlated

**(b)** Not significantly correlated

**Figure 4: Pairwise scatterplot with each point representing a single spectrum.** In (a), the peptides are identified as significantly correlated. In (b), the peptides are not significantly correlated. If the peptides in (a) share more than 25% of their ions, the lower scoring peptide (y-axis) will be removed.

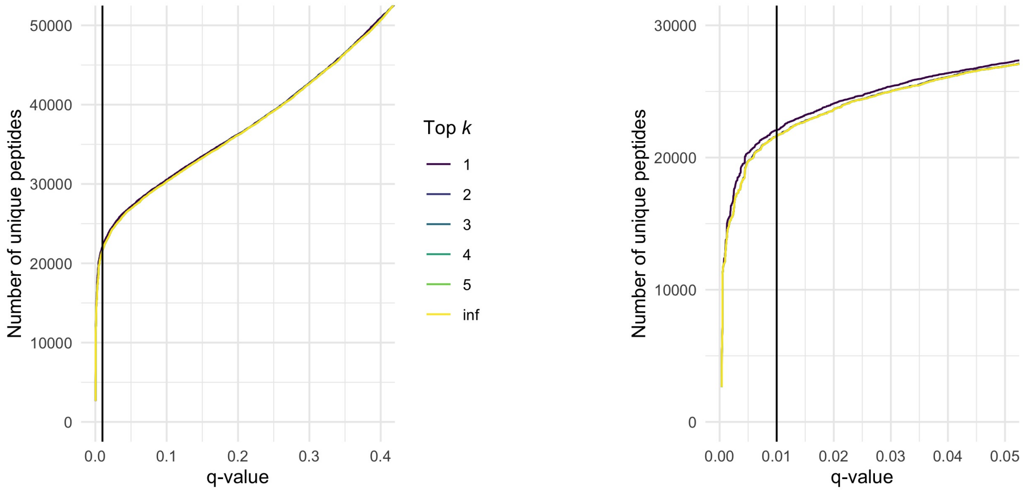


**(a)** q-value plot **(b)** Zoom in of (a)

**Figure 5: Plot of q-values when the top $k$ peptides are retained.** Here, the top $k$ scoring peptides for each spectrum are retained for FDR control. Although many DIA spectra are chimeric, it is rare that multiple ”true” peptides would share the same optimally matching spectrum. Therefore, using a $k$ value of 1 does not seem to eliminate a significant number of peptide matches, rather it increases power presumably by culling some high scoring fake matches. The vertical line represents 1% FDR threshold.

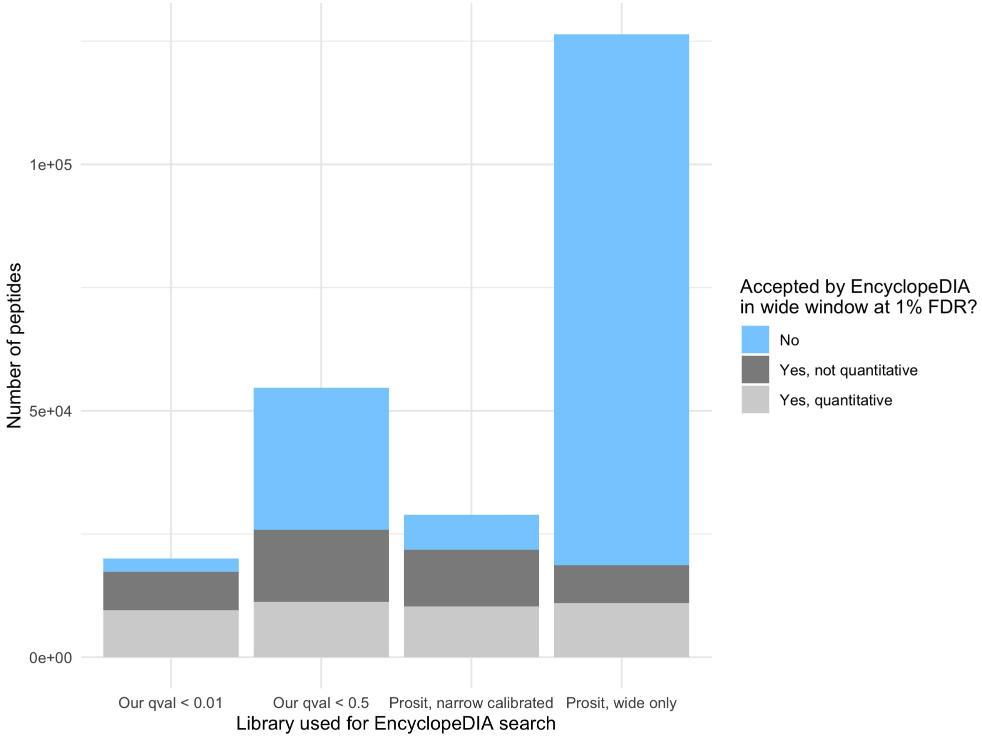


**Figure 6: Summary of the number of peptides in EncyclopeDIA runs using alternative libraries.** The figure summarizes the results of running EncyclopeDIA on the wide-window yeast DIA data using four different libraries: the first two runs used narrow-window yeast DIA data processed by our novel tool using a q-value cutoff of 0.01 and 0.5 to generate the library, and the third run used EncyclopeDIA’s ability to take advantage of the same narrow-window data to significantly reduce the initial Prosit-generated library. The last run used the full-size Prosit library of all possible tryptic peptides.
