## Supplemental File S2 for "A flexible workflow for building spectral libraries from narrow window data independent acquisition mass spectrometry data": precursor_matrix.py

```
#!/usr/bin/env python3
"""
This module creates a matrix, M, where the columns are scans, the
rows are precursors, and the values are the xcorr scores from
tide-search results for each DIA window.

You'll need the following prerequisite packages installed:
    tqdm
    numpy
    pandas

I've written this module such that it can either be run as script
from the command line, or such that the public functions (those without
a leading underscore) can be readily used in other python scripts. To
do so, you'll need to append this directory to your python path by
including the following in your script:

    import sys
    sys.path.append("path/to/here")
    import precursor_matrix

If using this within a python script, refer the docstrings and
type-hints should provide all of the information you need to know.
"""
import os
import time
import gzip
import logging
import argparse
from contextlib import ExitStack
from tempfile import TemporaryFile
from typing import Tuple, Dict

import numpy as np
import pandas as pd
from tqdm import tqdm  # For progress bars

# Public Functions ------------------------------------------------------------
def get_window(psm_file: str, window: str, out_file: str,
               force_: bool = False) -> str:
    """
    Extract the PSMs from a single window.

    Parameters
    ----------
    psm_file : str
        The search results for all windows.
    window : float
        The window, matching a precursor m/z.
    out_file: str
        The output file to write.

    Returns
    -------
    str
        The output file.
    """
    if os.path.isfile(out_file) and not force_:
        return out_file

    # handle gzipped files:
    if psm_file.endswith(".gz"):
        read_psms = gzip.open
    else:
        read_psms = open

    with read_psms(psm_file) as psms, open(out_file, "w+") as out:
        header = psms.readline()
        window_idx = header.split("\t").index("spectrum precursor m/z")

        out.write(header)
        for line in tqdm(psms, unit=" PSMs"):
            if line.split("\t")[window_idx] == window:
                out.write(line)

    return out_file


def construct(psm_file: str,
              output_dir: str = "windows-out",
              pickle_output: bool = False,
              score: str = "xcorr score",
              overwrite=False,
              liberal_matching=False) \
              -> Tuple[str]:
    """
    Construct a precursor by scan matrix of scores for each DIA window.

    The PSMs from a crux txt formatted file are converted to a wide
    matrix where the columns are scans, the rows a precursors and the
    value of each element is the assigned score. A matrix is created
    separately for each DIA window and individually saved in the
    specified output directory, either in a tab-delimited text format
    or as a gzipped, pickled Python object.

    Parameters
    ----------
    psm_file : str
        The PSMs from a tide-search in the crux txt format. If you want
        to include decoys, the search should be run with '--concat T'.

    output_dir : str
        The directory in which to save the matrix each DIA window.

    pickle_output : bool
        Store matrices as gzipped, pickled objects instead of tab-delmited text files.

    score : str
        The score column to use as values in matrices.

    overwrite : bool
        Overwrite existing result matrices.

    liberal_matching : bool
        If False, targets missing a corresponding decoy are discarded. If
        True, they are only discarded if the sequence between termini
        consists of only one amino acid.

    Returns
    -------
    window files
        A tuple containing the file corresponding to each DIA window.
    file mapping
        A string containing the file specifying how the MS data files map
        to indices.
    """
    os.makedirs(output_dir, exist_ok=True)

    # handle gzipped files:
    if psm_file.endswith(".gz"):
        read_psms = gzip.open
    else:
        read_psms = open

    # Set extension:
    if pickle_output:
        ext = ".pkl.gz"
    else:
        ext = ".txt"

    # STEP 1: Create temporary files containing the PSMs from each DIA window.
    window_temp_files = {}
    file_idx = 0
    file_mapping = {}
    logging.info("Sorting PSMs into DIA windows...")
    with ExitStack() as stack, read_psms(psm_file, "rt") as psms:
        # Find important columns:
        header = psms.readline()
        window_idx = header.split("\t").index("spectrum precursor m/z")
        file_col_idx = header.split("\t").index("file")

        # Write the line to the temp file if it exists.
        for line in tqdm(psms, unit=" PSMs"):
            line = line
            split_line = line.split("\t")
            win = split_line[window_idx]
            file_name = split_line[file_col_idx]

            if win in window_temp_files.keys():
                window_temp_files[win].write(line)
            else:
                window_temp_files[win] = stack.enter_context(TemporaryFile("w+t"))
                window_temp_files[win].write(header)
                window_temp_files[win].write(line)

            if not file_name in file_mapping.keys():
                file_mapping[file_name] = file_idx
                file_idx += 1

        logging.info("Mapped MS data files to indices:")
        file_names = []
        file_indices = []
        for file_name, file_idx in file_mapping.items():
            logging.info("\t%i = %s", file_idx, file_name)
            file_names.append(file_name)
            file_indices.append(file_idx)

        file_map_out = os.path.join(output_dir, "file_mapping.txt")
        pd.DataFrame({"index": file_indices, "file": file_names}) \
          .sort_values("index") \
          .to_csv(file_map_out, sep="\t", index=False)

        logging.info("Wrote file map to %s", file_map_out)

        # STEP 2: Load PSMs by window and transform to matrix.
        # Each window is saved as a separate file in 'out_dir'.
        logging.info("Transforming PSMs to matrices...")
        out_files = []
        unmapped_targets = 0
        mapped_targets = 0
        for window, fobj in tqdm(window_temp_files.items(), unit=" DIA windows"):
            out_file = os.path.join(output_dir, f"window_{int(float(window))}"+ext)
            out_files.append(out_file)

            # If the file is exists and we don't want to overwrite.
            if os.path.isfile(out_file) and not overwrite:
                logging.warning("%s already exist. Use 'overwrite=True' to "
                                "overwrite it.", out_file)
                continue

            # Create the matrix and save it.
            fobj.seek(0)
            df, unmapped, mapped = _process_window(fobj, score, liberal_matching,
                                                   file_mapping)
            unmapped_targets += unmapped
            mapped_targets += mapped
            if pickle_output:
                df.to_pickle(out_file)
            else:
                df.to_csv(out_file, sep="\t")

        if unmapped_targets:
            logging.warning("%i target precursors failed to be mapped to decoy "
                            "precursors and were discarded.", unmapped_targets)

        logging.info("%i target precursors were successfully mapped to decoy"
                     "precursors across all DIA windows.", mapped_targets)

    return tuple(out_files), file_mapping


# Utility Functions -----------------------------------------------------------
def _process_window(psm_file: str, score: str, liberal_matching: bool,
                    file_mapping: Dict[str, int]) \
    -> Tuple[pd.DataFrame, int, int]:
    """
    Read specific lines to a pandas DataFrame.

    This function reads specific rows of the tide-search results file and
    aggregates them into a matrix where the columns are scans, the rows
    are precursors, and the values are scores.

    Parameters
    ----------
    psm_file : str
        The tide-search.txt file.
    rows : array-like
        The indices of rows belonging to a DIA window.
    total_rows : int
        The number of lines in 'psm_file'
    score : str
        The name of the score column.
    liberal_matching : bool
        If False, targets missing a corresponding decoy are discarded. If
        True, they are only discarded if the sequence between termini
        consists of only one amino acid.
    file_mapping : Dict mapping str to int
        The file names and their corresponding indices.

    Returns
    -------
    pandas.DataFrame
        The PSMs
    int
        The number of unmatched target precursors
    int
        The number of matched target precursors.
    """
    cols = ("file","scan", "sequence", "charge", "target/decoy",
            "original target sequence", score)

    col_types = {"file": "str",
                 "scan": "str",
                 "sequence": "str",
                 "charge": "str",
                 "target/decoy": "str",
                 "original target sequence": "str",
                 score: np.float64}

    df = pd.read_csv(psm_file, sep="\t", usecols=cols,
                     dtype=col_types, engine="c")

    # Create a unique precursor id and set it as the index.
    df["file"] = df["file"].replace(file_mapping).astype(str)
    df["precursor"], unmapped, mapped = _pair_decoys(df, liberal_matching)

    if not liberal_matching:
        df = df.loc[df["precursor"].str.endswith("_"), :]

    # dropna() fails when dataframes are too big:
    # https://github.com/pandas-dev/pandas/issues/35227
    #df = df.dropna()
    df = df.loc[(~df.isna()).product(axis=1).astype(bool), :]

    df["precursor"] = df["file"] + "_" + df["precursor"]
    df = df.set_index("precursor").rename(columns={score: "score"})

    # Keep only relevant columns:
    df = df.loc[:, ["score", "scan"]]
    logging.debug("The first row of 'df' is:\n%s", df.iloc[0, :])

    # Pivot to wide DataFrame.
    try:
        df = df.pivot(columns="scan", values="score")
    except ValueError:
        dups = df.index[df.index.duplicated()].tolist()
        logging.debug("The first duplicate is: %s", str(dups[0]))
        dups = "\n\t".join(dups)
        raise ValueError(f"Duplicate precursors found in window:\n\t{dups}")

    df.index.name = None
    df.columns.name = None

    return df, unmapped, mapped


def _pair_decoys(df: pd.DataFrame, liberal_matching: bool) \
    -> Tuple[pd.Series, int, int]:
    """
    Build a mapping between target sequences and their corresponding decoys.

    Parameters
    ----------
    df : pandas.DataFrame
        A dataframe of the parsed PSMs.

    liberal_matching : bool
        If False, targets missing a corresponding decoy are discarded. If
        True, they are only discarded if the sequence between termini
        consists of only one amino acid.

    Returns
    -------
    tuple of pd.Series, int, and int
        A unique precursor identifier, where decoys are mapped to their
        corresponding targets, the number of unmatched target precursors
        and the number of matched target precursors.
    """
    cols = ["original target sequence", "charge"]
    df["target"] = df["target/decoy"] == "target"
    prec = df.loc[:, ["target", "sequence"] + cols].drop_duplicates()

    # Add a row for the case when no peptides have modifications.
    prec = prec.append({"target": True,
                        "sequence": "X[X]",
                        "original target sequence": np.nan,
                        "charge": np.nan},
                       ignore_index=True)

    # All modifications are sorted and pasted together.
    # This is the key to matching modified targets to their modified decoys.
    prec["mods"] = (prec.sequence.str.extractall("(.\[.+?\])")
                                 .sort_values(by=0)
                                 .groupby(level=0)
                                 .apply(lambda x: "".join(x[0])))

    prec = prec.fillna("")
    decoy_map = {}
    unmapped_targets = []
    for mod, pep_group in prec.groupby(["mods"] + cols):
        targets = pep_group[pep_group.target].sample(frac=1)
        decoys = pep_group[~pep_group.target].sample(frac=1)

        for idx, decoy in enumerate(decoys.sequence):
            # For finite k, a decoy may be scored when its target is not.
            if idx < len(targets):
                decoy_map[decoy] = targets.sequence.iloc[idx]
            else:
                decoy_map[decoy] = decoy

        # Handle specific cases cases:
        # 1. There are more decoys than targets for the same sequence.
        if len(decoys) and idx+1 < len(targets) and not liberal_matching:
            unmapped_targets += targets.sequence.iloc[idx+1:].tolist()

        # 2. There are no decoys
        elif not len(decoys) and not liberal_matching:
            unmapped_targets += targets.sequence.tolist()

        # 3. There are no decoys, but you want to be lenient.
        # In this case, only precursors in which all of the AA's between
        # termini are identical are discarded.
        elif not len(decoys) and liberal_matching:
            mid_seq = (targets.sequence.str.extract("^.(.+).$", expand=False)
                                       .apply(lambda x: len(set(x))))

            for seq in mid_seq:
                if seq == 1:
                    unmapped_targets += targets.sequence.tolist()

    # Do the assignments
    unmapped_targets = set(unmapped_targets)
    df["match_sequence"] = df.apply(_replace_decoy, axis=1, decoy_dict=decoy_map,
                                    unmapped_targets=unmapped_targets)

    df["td"] = "D"
    df.loc[df.target, "td"] = "T"
    precursor = df.td + "_" + df.sequence + "_" + df.charge + "_" + df.match_sequence

    na_precursor = pd.isna(precursor)
    if pd.isna(precursor).sum():
        df.loc[na_precursor].to_csv("error.csv")
        raise ValueError("NA values detected. See 'error.csv'.")

    return precursor, len(unmapped_targets), len(decoy_map)


def _replace_decoy(df, decoy_dict, unmapped_targets) -> str:
    """
    If the PSM is a decoy, replace the decoy with its corresponding
    target

    Parameters
    ----------
    df : pd.DataFrame
        A 1 row dataframe containing the precursor identifying information.

    decoy_dict : dict
        A mapping of decoy sequences to their corresponding target.

    unmapped_targets : set
        The set of target sequences that don't have a corresponding decoy.

    Returns
    -------
    str
        The replaced sequence
    """
    new_seq = df.sequence
    if df.target and df.sequence in unmapped_targets:
        new_seq = ""
    elif not df.target:
        new_seq = decoy_dict[df.sequence]

    return new_seq


# MAIN ------------------------------------------------------------------------
def main():
    """This is executed when run as a command line program."""
    st_time = time.time()

    desc = """
    Construct a precursor by scan matrix of scores for each DIA window.

    The PSMs from a crux txt formatted file are converted to a wide
    matrix where the columns are scans, the rows a precursors and the
    value of each element is the assigned score. A matrix is created
    separately for each DIA window and individually saved in the
    specified output directory, either in a tab-delimited text format
    or as a gzipped, pickled Python object.
    """

    parser = argparse.ArgumentParser(description=desc)
    parser.add_argument("psm_file", type=str,
                        help=("The PSMs from a tide-search in the crux txt "
                              "format. If you want to include decoys, the "
                              "search should be run with '--concat T'."))

    parser.add_argument("--output_dir", type=str, default="window-out",
                        help=("The directory in which to save the matrix for "
                              "each DIA window."))

    parser.add_argument("--pickle_output", default=False, action="store_true",
                        help=("Store matrices as gzipped, pickled Python "
                              "objects instead of tab-delmited text files."))

    parser.add_argument("--score", type=str, default="xcorr score",
                        help=("The score column to use as values in the "
                              "matrices"))

    parser.add_argument("--overwrite", default=False, action="store_true",
                        help=("Overwrite existing result matrices."))

    parser.add_argument("--seed", type=int, default=42,
                        help=("The random seed. Some targets must be randomly "
                              "paired with their corresponding decoy. Setting "
                              "the seed ensures reproducibility."))

    parser.add_argument("--liberal_matching", default=False, action="store_true",
                        help=("Allow less stringent assumptions for matching "
                              "target and decoy precursors."))

    parser.add_argument("--debug", default=False, action="store_true",
                        help=("Print debugging messages."))

    args = parser.parse_args()

    if args.debug:
        level = logging.DEBUG
    else:
        level = logging.INFO

    logging.basicConfig(level=level, format="%(levelname)s: %(message)s")

    np.random.seed(args.seed)
    fout = construct(psm_file=args.psm_file,
                     output_dir=args.output_dir,
                     pickle_output=args.pickle_output,
                     score=args.score,
                     overwrite=args.overwrite,
                     liberal_matching=args.liberal_matching)

    logging.info("Wrote matrices for %i DIA windows to %s.", len(fout),
                 args.output_dir)

    end_time = time.time()
    logging.info("Completed in %0.2f min.", (end_time-st_time)/60)

if __name__ == "__main__":
    main()
```
