## Supplemental File S3 for "A flexible workflow for building spectral libraries from narrow window data independent acquisition mass spectrometry data": process_precursors.R

```
#!/usr/bin/env Rscript
suppressPackageStartupMessages(library(tidyverse, quietly = T))
suppressPackageStartupMessages(library(smoother, quietly = T))
suppressPackageStartupMessages(library(changepoint, quietly = T))
suppressPackageStartupMessages(library(tictoc, quietly = T))
suppressPackageStartupMessages(library(reshape2))
suppressPackageStartupMessages(library(magrittr))
suppressPackageStartupMessages(library(reticulate))

#use_condaenv("updated_base")
peptides <- import_from_path("peptides", path = "bin/")

source_python("bin/pepsim_forR.py")

# Functions --------------------------------------------------------------------

#' Print the help message.
print_help <- function() {
  msg <- c("USAGE: peptide_groups.R  [--num_scans <int>] [--matrix_dir <dir>] <matrices>",
           "",
           "Preprocess the matrices for between 'precursor_matrix.py' and ",
           "'precursor_confidence.py. The output files are written to the ",
           "directory written to the current working directory, or to the ",
           "directory specified by '--matrix_dir",
           "",
           "Arguments:",
           "    matrices   - One or more matrix from 'precursor_matrix.py",
           "    num_scans - Number of spectra to include in acos grouping. If no number, all spectra will be used",
           "    matrix_dir - The directory in which to look for 'matrices'",
           "",
           "Output:",
           "    processed_matrices - Matrices with adjusted scores",
           "    file_names         - A list of the processed matrix files",
           "    width_map          - A mapping of peptides to the selected width",
           "")

  return(cat(msg, sep = "\n"))
}

#' Parse the number of scans from command line
#' This probably isn't how I should to this, but it seems to work
#'
#' @return A value indicating number of scans to use later on.

parse_scans <- function() {
  args <- commandArgs(trailingOnly = T)

  if ("-h" %in% args | "--help" %in% args) {
    print_help()
    quit()
  }

  # Parse matrix_dir, if provided:
  
  if (args[1] == "--num_scans") {
    num_scans <- args[2]
    args <- args[3:length(args)]
  } else {
    num_scans <- "all"
  }
  return(num_scans)
}


#' Parse the command line arguments
#'
#' This function does support globbing (using the *). Also, if no files are
#' provided, it will assume you want any file in your working directory that
#' ends with '.txt'
#'
#' @return A vector containing the matrix_dir followed by the matrix filenames.
parse_args <- function() {
  args <- commandArgs(trailingOnly = T)

  if ("-h" %in% args | "--help" %in% args) {
    print_help()
    quit()
  }

  if (args[1] == "--num_scans") {
    num_scans <- args[2]
    args <- args[3:length(args)]
  } else {
    num_scans <- "all"
  }
  
  # Parse matrix_dir, if provided:
  if (args[1] == "--matrix_dir") {
    matrix_dir <- args[2]
    args <- args[3:length(args)]
  } else {
    matrix_dir <- "."
  }

  # Find the files:
  if (length(args) == 0) {
    files <- list.files(pattern = "*.txt$")
  } else {
    files <- c()
    for (filepattern in args) {
      filepattern <- paste0(filepattern, "$")
      found_files <- list.files(path = matrix_dir,
                                pattern = filepattern,
                                full.names = T)

      files <- c(files, found_files)
    }
  }

  if (length(files) == 0) {
    cat("ERROR: No matrix files were found!\n")
    quit(status = 1)
  }

  return(c(matrix_dir, files))
}


#' Parse a precursor matrix into a DataFrame.
#'
#' @param matrix_file The matrix file to parse.
#' @return A DataFrame, where columns are scans and rows are precursors.
parse_matrix <- function(matrix_file) {
  cat(paste0("INFO: Reading in ", matrix_file, " :)\n"))

  mat <- read.delim(matrix_file, check.names = FALSE, row.names = 1)
  mat <- mat[ , order(as.numeric(names(mat)))]
  cat(paste0("INFO: ", matrix_file, " has ", nrow(mat), " precursors and ", ncol(mat), " spectra :)\n"))

  if (sum(is.na(mat)) > 0) {
    stop(paste0("Missing values encountered in ", matrix_file))
  }
  return(mat)
  
}


#' Detect changepoint
#'
#' @param mat The matrix returned from parse_matrix
#' @return a shortened dataframe
filter_scans <- function(mat){
  temp <- data.frame(median = apply(mat, 2, median)) %>%
          mutate(spectrum = as.numeric(colnames(mat))) %>%
          arrange(spectrum)
  change <- cpt.meanvar(temp$median, method = "BinSeg", Q = 2)
  low <- temp[seg.len(change)[1], 2] %>%
         as.numeric()
  high <- temp[seg.len(change)[1]+seg.len(change)[2], 2] %>%
          as.numeric()

  if ((high-low) <= (0.5*as.numeric(colnames(mat[ncol(mat)])))) {
    cat("WARNING: More than 50% of spectra were excluded, repeating with 3 changepoints \n")

    change <- cpt.meanvar(temp$median, method = "BinSeg", Q = 3)
    low <- temp[seg.len(change)[1], 2] %>%
           as.numeric()
    high <- temp[seg.len(change)[1]+seg.len(change)[2]+seg.len(change)[3], 2] %>%
            as.numeric()
  }

  if ((high-low) <= (0.5*as.numeric(colnames(mat[ncol(mat)])))) {
    cat("WARNING: More than 50% of spectra were excluded, repeating with 4 changepoints \n")

    change <- cpt.meanvar(temp$median, method = "BinSeg", Q = 4)
    low <- temp[seg.len(change)[1]+seg.len(change)[2], 2] %>%
           as.numeric()
    high <- temp[seg.len(change)[1]+seg.len(change)[2]+seg.len(change)[3]+seg.len(change)[4], 2] %>%
            as.numeric()
  }

  if ((high-low) <= (0.5*as.numeric(colnames(mat[ncol(mat)])))) {
    cat("WARNING: More than 50% of spectra were excluded with 4 changepoints, check this file! \n")
  }

  cat(paste0("INFO: Retaining spectra from  ", low, " to ", high, " :)\n"))
  mat <- select(mat, num_range("", low:high))

  return(mat)
}


#' Determine peak widths based on relative RZ score
#'
#' @param mat The shortened matrix returned from filter_scan
#' @return a long dataframe containing:
#'      sequence: <file index>_<target or decoy>_<precursor sequence>_<charge state>_<original target sequence>
#'      peak width (width)
#'      distance from max to left peak boundary (left_id)
#'      distance from max to right peak boundary (right_id)
assign_width <- function(mat){
  norm_coeff <- apply(mat, 1, median) %>%
    data.frame() %>%
    cbind(MAD = apply(mat, 1, mad))

  mat_temp <- cbind(
    data.frame(matrix(0, ncol = 20, nrow = nrow(mat))),
    mat,
    data.frame(matrix(0, ncol = 20, nrow = nrow(mat)))
  ) %>%
    t() %>%
    data.frame()

  mat_temp <- mapply('-', mat_temp, norm_coeff[,1]) %>%
    data.frame(check.names = FALSE)

  mat_temp <- mapply('/', mat_temp, norm_coeff[,2]) %>%
    t() %>%
    data.frame(check.names = FALSE)

  row.names(mat_temp) <- row.names(mat)
  colnames(mat_temp)[21:(ncol(mat)+20)] <- colnames(mat)

  meta <- bind_cols(
    sequence = row.names(mat_temp),
    top_scan_idx = apply(mat_temp, 1, which.max),
    top_scan = colnames(mat_temp)[apply(mat_temp,1,which.max)],
    top_rzscore = apply(mat_temp, 1, max)
  )

  max_df <- data.frame(
    mat_temp[1, (meta$top_scan_idx[1]-20):(meta$top_scan_idx[1]+20)]
  ) %>%
    set_colnames(c(seq(1:41))) %>%
    bind_cols(sequence = (meta$sequence[1]))

  for (x in 2:nrow(mat_temp)) {
    max_x <- data.frame(
      mat_temp[x, (meta$top_scan_idx[x]-20):(meta$top_scan_idx[x]+20)]
    ) %>%
      set_colnames(c(seq(1:41))) %>%
      bind_cols(sequence = (meta$sequence[x]))

    max_df <- bind_rows(max_df, max_x)
  }

  rm(mat_temp)

  max_df_rel <- cbind(max_df, max_score = meta$top_rzscore) %>%
                melt(id.vars = c("sequence", "max_score"),
                  value.name = "raw_score") %>%
                mutate(seq = gsub(".*_", "", sequence)) %>%
                group_by(seq) %>%
                top_n(n = 1, wt = max_score) %>%
                top_n(n = 1, wt = raw_score) %>%
                select(seq, max_score)


  meta <- melt(max_df, id.vars = c("sequence"),
          value.name = "raw_score", variable.name = "scan") %>%
          mutate(seq = gsub(".*_", "", sequence)) %>%
          full_join(max_df_rel, by = "seq") %>%
          mutate(rel_score = raw_score/max_score) %>%
          mutate(scan = as.numeric(as.character(scan))) %>%
          mutate(col_id = (scan-21)) %>%
          mutate(binary_score = if_else(rel_score >= 0.75, 1, 0)) %>%
          mutate(binary_score = if_else(col_id == 0, 1, binary_score)) %>%
          group_by(sequence, scan) %>%
          slice(1) %>%
          ungroup() %>%
          arrange(sequence, scan) %>%
          group_by(sequence, grp = with(rle(binary_score),
            rep(seq_along(lengths), lengths))) %>%
          mutate(Counter = seq_along(grp)) %>%
          ungroup() %>%
          inner_join(select(filter(., col_id == 0), grp), by = "grp") %>%
          group_by(sequence) %>%
          arrange(-Counter) %>%
          mutate(left_id = min(col_id)) %>%
          mutate(right_id = max(col_id)) %>%
          mutate(width_1 = max(as.numeric(Counter))) %>%
          slice(1) %>%
          group_by(seq) %>%
          mutate(width = max(width_1)) %>%
          mutate(left_id = if_else(width_1 == width, left_id,
            if_else(width %%2 == 0, -(width/2) + 1, -(width-1)/2 ))) %>%
          mutate(right_id = if_else(width_1 == width, right_id,
            if_else(width %%2 == 0, (width/2), (width-1)/2 ))) %>%
          ungroup() %>%
          select(sequence, width, left_id, right_id) %>%
          full_join(meta, by = "sequence") %>%
          mutate(left = top_scan_idx + left_id) %>%
          mutate(right = top_scan_idx + right_id) %>%
          select(sequence, width, left_id, right_id)
          
  cat("INFO: Selected window widths\n")

  return(meta)
}


#' Row-wise Tailor normalization
#'
#' @param mat The matrix returned from filter_scans
#' @return a matrix with row normalized scores
row_norm <- function(mat){
  #norm_mat <- data.frame(t(mat))

  norm_coeff <- apply(mat, 1, quantile, probs = c(0.99)) #%>%
                #data.frame()
                
    #norm_mat <- mapply('/', norm_mat, norm_coeff[,1]) %>%
  #            t() %>%
   #           data.frame(check.names = FALSE)
   
   norm_mat <- data.frame(mat/norm_coeff)

  rownames(norm_mat) <- rownames(mat)
  colnames(norm_mat) <- colnames(mat)

  return(norm_mat)
  
}


#' Combine information about each precursor
#'
#' @param norm_mat, Tailor normalized scores from row_norm
#' @param meta, long data frame from assign_width
#' @return a dataframe with metadata about each precursor. Columns are:
#'      sequence: <file index>_<target or decoy>_<precursor sequence>_<charge state>_<original target sequence>
#'      score: maximum row-normalized score for that precursor
#'      top_scan: scan number of top score for precursor
#'      top_scan_idx: column index of top score for precursor, refers to mat and norm_mat
#'      width: assigned peak width (in cycles or columns)
#'      left: spectrum number of left peak edge
#'      right: spectrum number of right peak edge
#'      type: T or D to indicate target or decoy
#'      keep: variable we assign later, everything starts with 0
extract_meta <- function(norm_mat, meta){
  output_df <- bind_cols(
    sequence = row.names(norm_mat),
    score = data.frame(apply((norm_mat), 1, max)),
    top_scan = data.frame(apply(norm_mat, 1, which.max)),
    top_scan_idx = data.frame(apply(norm_mat, 1, which.max))
  ) %>%
    set_colnames(c("sequence", "score", "top_scan", "top_scan_idx"))  %>%
    mutate(
      top_scan=as.numeric(gsub("\\..*", "", colnames(norm_mat[top_scan])))
    ) %>%
    full_join(meta, by = "sequence") %>%
    mutate(left = top_scan_idx+left_id) %>%
    mutate(left = if_else(left <= 0, 1, left)) %>%
    mutate(left = colnames(norm_mat)[left]) %>%
    mutate(right = top_scan_idx+right_id) %>%
    mutate(right = if_else(right >= ncol(norm_mat), 1, right)) %>%
    mutate(right = colnames(norm_mat)[right]) %>%
    mutate(seq = gsub(".*_", "", sequence)) %>%
    mutate(type = unlist(strsplit(sequence, "_"))[seq(from=2, by=5, length.out = nrow(meta))]) %>%
    arrange(desc(score)) %>%
    group_by(seq) %>%
    slice(1)  %>%
    mutate(keep = 0) %>%
    ungroup() %>%
    #group_by(top_scan) %>%
    #mutate(rank = rank(-score)) %>%
    #ungroup() %>%
    select(-left_id, -right_id, -seq)

  return(output_df)
}


#' Define angle cosine function
#'
#' @param x, a vector of scores ordered by time
#' @param y, a vector of scores ordered by time
#' @return an angle cosine value
angle <- function(x,y){
  dot.prod <- x%*%y
  norm.x <- sqrt(rowSums(x^2))
  norm.y <- sqrt(colSums(y^2))
  theta <- (dot.prod / (norm.x * norm.y))
  as.numeric(theta)
}


#' Remove highly correlated precursors
#'
#' @param TD, either "T" or "D" if targets or decoys are considered
#' @param output_df, output from extract_meta
#' @param mat, un-normalized scores from filter_scans
#' @param num_scans, Number of spectra to include in acos grouping
#' @return a dataframe with metadata about each precursor:
#'      Rank: rank of score of precursor relative to all others, 1 = highest score
#'      sequence: <file index>_<target or decoy>_<precursor sequence>_<charge state>_<original target sequence>
#'      score: maximum row-normalized score for that precursor
#'      top_scan: scan number of top score for precursor
#'      top_scan_idx: column index of top score for precursor, refers to mat and norm_mat
#'      width: assigned peak width (in cycles or columns)
#'      left: spectrum number of left peak edge
#'      right: spectrum number of right peak edge
#'      type: T or D to indicate target or decoy
#'      keep: variable indicating whether we will keep precursor (1) ir exclude as a hitchhiker (2)
acos_group <- function(TD, output_df, mat, num_scans){
  
  meta_full_target <- output_df %>%
    filter(type == "T") %>%
    arrange(desc(score)) %>%
    tibble::rownames_to_column(var = "Rank") %>%
    mutate(Rank = as.numeric(Rank)) %>%
    mutate(cor_with = "NONE")

  meta_full_decoy <- output_df %>%
    filter(type == "D") %>%
    arrange(desc(score)) %>%
    tibble::rownames_to_column(var = "Rank") %>%
    mutate(Rank = as.numeric(Rank)) %>%
    mutate(cor_with = "NONE")

  mat_target <- t(mat) %>%
    data.frame() %>%
    set_colnames(row.names(mat)) %>%
    select(meta_full_target$sequence)

  mat_decoy <- t(mat) %>%
    data.frame() %>%
    set_colnames(row.names(mat)) %>%
    select(meta_full_decoy$sequence)

  if (TD == "T"){
    meta_full_check <- meta_full_target
    mat_check <- mat_target
    mat_null <- mat_decoy
  }

  if (TD == "D"){
    meta_full_check <- meta_full_decoy
    mat_check <- mat_decoy
    mat_null <- mat_target
  }
  
  num_scans_2 <- num_scans
  
  if (num_scans_2 == "all"){
      num_scans_2 <- nrow(mat_check)
  }
  
  num_scans_2 <- as.integer(num_scans_2)
  
  if (num_scans_2 > nrow(mat_check)){
      num_scans_2 <- nrow(mat_check)
  }
  
  cat(paste0("INFO: Using top ", num_scans_2, " spectra for acos grouping.\n"))

  for (k in 1:nrow(meta_full_check)){
    
    precursor <- meta_full_check[k,]
    
    selected <- select(mat_check, precursor$sequence)
        
    precursor_cols <- selected %>%
      bind_cols(spectrum = row.names(.)) %>%
      bind_cols(index = row.names(.)) %>%
      arrange(desc(.[,1])) %>%
      slice(1:num_scans_2) %>%
      select(spectrum) %>%
      unlist()
      
    selected <- as.matrix(selected[precursor_cols,])
            
    mat_check_2 <- mat_check[precursor_cols,]
    
    mat_null_2 <- mat_null[precursor_cols,]


    if(precursor$keep != 2) {
      meta_full_check$keep[k] <- 1
    }

    shared_window <- filter(
      meta_full_check,
      ((left <= meta_full_check$right[k] & left >= meta_full_check$left[k]) |
       (right <= meta_full_check$right[k] & right >= meta_full_check$left[k]) |
       (left<=meta_full_check$top_scan[k] & right >= meta_full_check$top_scan[k])|
       ( meta_full_check$left[k] <= top_scan & meta_full_check$right[k] >= top_scan))
    ) %>%
      filter(sequence != precursor$sequence) %>%
      filter(Rank > precursor$Rank) %>%
      filter(keep != 2)

    if(nrow(shared_window) != 0) {
      sample_subset <- select(
        mat_null_2,
        sample(
          seq(1, ncol(mat_null_2), 1),
          size = if_else(ncol(mat_null_2) > 1000, 1000, as.numeric(ncol(mat_null_2))),
          replace = FALSE
        )
      )

      #selected <- as.matrix(select(mat_check, precursor$sequence))
      sample_subset <- angle(t(as.matrix(sample_subset)), selected) %>%
        data.frame() %>%
        set_colnames("acos") %>%
        summarise(max = max(acos))

      shared_mat <- as.matrix(mat_check_2[shared_window$sequence])
      shared_check <- angle(t(shared_mat), selected) %>%
        data.frame() %>%
        set_colnames("acos") %>%
        mutate("keep" = if_else(acos <= sample_subset$max, 1, 2)) %>%
        mutate(cor_with = if_else(acos <= sample_subset$max, "NONE", precursor$sequence))
        #mutate("keep" = (acos <= sample_subset$max) + 1)

      meta_full_check[shared_window$Rank, "keep"] <- shared_check$keep
      
      meta_full_check[shared_window$Rank, "cor_with"] <- shared_check$cor_with
    }
  }
  
  all_sequences <- meta_full_check %>%
    mutate(seq = gsub(".*_","",sequence)) %>%
    mutate(seq_with = gsub(".*_","",cor_with)) %>%
    mutate(mz = 100) %>%
    filter(keep == 2) %>%
    mutate(cor = 0)
  
  for(l in 1:nrow(all_sequences)){
    cor <- calc_proportion_fragments_incommon(all_sequences$seq[l], all_sequences$seq_with[l], 0.02)
    if (cor < 0.25){
        meta_full_check$keep[all_sequences$Rank[l]] <- 1
    }
  }
  
  return(meta_full_check)
}


#' Gather all precursors that will be kept together
#'
#' @param meta_full_target, output from acos_group using TD = "T"
#' @param meta_full_decoy, output from acos_group using TD = "D"
#' @param output_df, output from extract_meta
#' @returns information on all targets and decoys that will be kept based on this procedure:
#'      sequence: <file index>_<target or decoy>_<precursor sequence>_<charge state>_<original target sequence>
#'      score: maximum row-normalized score for that precursor
#'      top_scan: scan number of top score for precursor
#'      top_scan_idx: column index of top score for precursor, refers to mat and norm_mat
#'      width: assigned peak width (in cycles or columns)
#'      left: spectrum number of left peak edge
#'      right: spectrum number of right peak edge
#'      type: T or D to indicate target or decoy
combine_meta <- function(meta_full_target, meta_full_decoy, output_df){
  summary_meta <- rbind(meta_full_target, meta_full_decoy) %>%
                  filter(keep != 2) %>%
                  select(-keep)

  cat(paste0("INFO: Excluded ", nrow(filter(meta_full_target, keep == 2)),
        " targets and ", nrow(filter(meta_full_decoy, keep == 2)),
        " decoys due to high correlation.\n"))

  return(summary_meta)
}


#' Save all the widths
#'
#' @param summary_meta, output from combine_meta
#' @return width_map for this window with column for sequence (precursor identifier) and width
extract_width <- function(summary_meta){
    width_map <- select(summary_meta, sequence, width)

  return(width_map)
}


#' Make a wide matrix of remaining precursors
#'
#' @param summary_meta, output from combine_meta
#' @return output_df wide matrix of scores for window
new_matrix <- function(summary_meta){
    output_mat <- select(summary_meta, score, top_scan, sequence) %>%
               tidyr::spread(key=top_scan, value = score) %>%
               data.frame(check.names = FALSE)

  rownames(output_mat) <- output_mat$sequence
  output_mat <- subset(output_mat, , -c(sequence))

  return(output_mat)
}

# MAIN -------------------------------------------------------------------------

#' The main function
#'
#' This is what is run when the script is used from the command line.
main <- function() {
  num_scans <- parse_scans()
  files <- parse_args()
  matrix_dir <- files[1]
  files <- files[2:length(files)]
  filenames <- data.frame(col = " ")
  width_map <- data.frame(sequence = character(),
                width = integer())

  # Loop through files, applying the pipeline:
  for (matrix_file in files) {
    
    tic("TIME: Read in file")
    mat <- parse_matrix(matrix_file)
    toc(log = TRUE)
    
    tic("TIME: Changepoint detection")
    mat <- filter_scans(mat)
    toc(log = TRUE)
    
    tic("TIME: Assigned width")
    meta <- assign_width(mat)
    toc(log = TRUE)
    
    tic("TIME: Normalized scores")
    norm_mat <- row_norm(mat)
    toc(log = TRUE)
    
    tic("TIME: Extracted meta data")
    output_df <- extract_meta(norm_mat, meta)
    toc(log = TRUE)

    tic("TIME: Acos grouping on targets")
    meta_full_target <- acos_group("T", output_df, mat, num_scans)
    toc(log = TRUE)

    tic("TIME: Acos grouping on decoys")
    meta_full_decoy <- acos_group("D", output_df, mat, num_scans)
    toc(log = TRUE)

    tic("TIME: Concatenate acos data")
    summary_meta <- combine_meta(meta_full_target, meta_full_decoy, output_df)
    toc(log = TRUE)

    tic("TIME: Generate width map")
    width_map_sub <- extract_width(summary_meta)
    toc(log = TRUE)

    tic("TIME: Spread matrix")
    output_mat <- new_matrix(summary_meta)
    toc(log = TRUE)
    
    toc(log = TRUE)
    
    tic.log(format = TRUE)

    width_map <- bind_rows(width_map, width_map_sub)
    filenames <- cbind(filenames, col =
                    paste0(matrix_dir, "/processed_", sub(".*/", "", matrix_file)))
    
    write.table(output_mat, file =
                paste0(matrix_dir, "/processed_", sub(".*/", "", matrix_file)),
                sep = "\t", quote = FALSE, na = "")
      }
  
  width_map <- width_map %>%
                   tibble::column_to_rownames(var = "sequence")

  write.table(width_map, file = paste0(matrix_dir, "/processed_widths.txt"), quote = FALSE, sep = "\t")
  
  write.table(filenames, file = paste0(matrix_dir, "/processed_filenames.txt"), sep = " ",
        quote = FALSE, col.names = FALSE, row.names = FALSE)
  
  #print(head(mat_df))
}

# Only run from the main from the command line:
if (!interactive()) {
  main()
}
```
