## Supplemental File S1 for "A flexible workflow for building spectral libraries from narrow window data independent acquisition mass spectrometry data": precursor_confidence.py

```
#!/usr/bin/env python3
"""
This module estimate the FDR for detected precursors using the
spectrum-centric approach, the peptide-centric approach and
interpolating in-between.

You'll need the following prerequisite packages installed:
    tqdm
    numba
    numpy
    pandas

I've written this module such that it can either be run as script
from the command line, or such that the public functions (those without
a leading underscore) can be readily used in other python scripts. To
do so, you'll need to append this directory to your python path by
including the following in your script:

    import sys
    sys.path.append("path/to/here")
    import precursor_confidence.py

If using this within a python script, refer the docstrings and
type-hints should provide all of the information you need to know.
"""
import os
import glob
import time
import gzip
import logging
import argparse
from contextlib import ExitStack
from tempfile import TemporaryFile
from typing import Tuple, Union, Dict, Optional

import numba as nb     # For fast q-value calculations
import numpy as np
import pandas as pd
from tqdm import tqdm  # For progress bars

# Public Functions ------------------------------------------------------------
def assign(psm_matrix: Union[str, Tuple[str], pd.DataFrame, Tuple[pd.DataFrame]],
           topk: Union[int, Tuple[int], None] = 1,
           plus_one: bool = False,
           desc: bool = True,
           groups: Optional[Union[str, pd.DataFrame]] = None) -> pd.DataFrame:
    """
    Estimate q-values using the top-k matches per spectrum.

    When k = 1, this is equivalent to the spectrum-centric search. When
    k = inf, this is equivalent peptide-centric search. Practically, the
    maximum value of k will not exceed the value used for the 'top-match'
    parameter during the tide-search run.

    The format of the 'psm_matrix'---which can be provided either as the
    a file name or a pandas dataframe---should be such that rows are
    precursors, columns are scans, and values are the score. Furthermore,
    the precursor index should be of the format <T|D>_<sequence>_<charge>
    because this string is parsed to determine if the precursor was a
    target or decoy hit. Matrices of this type are saved by the
    'precursor_matrix.py' script.

    Parameters
    ----------
    psm_matrix : str, pandas.DataFrame or tuple of either
        One or more matrix in the format specified above. These can
        either be provided as one or more files to be parsed, or
        as one or more pandas.DataFrame.

    topk : int or tuple of int or None
        One or more value of k to use. k=Inf is always performed.
        If None, only k=Inf is performed.

    plus_one : bool
        Should the conservative addition of 1 to the numerator be used
        during FDR estimation?

    desc : bool
        Are higher scores better?

    groups : str or pandas.DataFrame
        A tab-delimited file or DataFrame with two columns to specify
        groups for group FDR. The first column should be the precursor,
        and the second should be the group. If None, no grouping is
        used.

    Returns
    -------
    pandas.DataFrame
        A DataFrame where rows are the target precursors and columns
        are the q-values for the indicated value of 'k'.
    """
    if isinstance(psm_matrix, str) or isinstance(psm_matrix, pd.DataFrame):
        psm_matrix = (psm_matrix, )

    if isinstance(topk, int):
        topk = np.array([topk, np.inf])
    elif topk is None:
        topk = np.array([np.inf])
    else:
        topk = np.array(list(topk) + [np.inf])

    # Sort topk so that higher values are first.
    topk = np.flip(np.sort(topk))

    # Read PSM data matrices
    logging.info("Reading and preprocessing PSM data matrices...")
    df = pd.concat([_parse_matrix(m, desc=desc, window_index=i)
                    for i, m in enumerate(tqdm(psm_matrix, unit="matrix"))])

    # Get grouping information
    if isinstance(groups, str):
        groups = pd.read_csv(groups, sep="\t").reset_index()
    elif groups is None:
        groups = pd.DataFrame({"key": df.key.unique(),
                               "group": 0})

    groups.columns = ["key", "group"]
    df = pd.merge(df, groups, how="left")

    # Estimate q-values for each value of k.
    logging.info("Estimating q-values for %i values of k...", len(topk))
    results = []
    prec_cols = ["score", "target", "precursor"]
    for k in tqdm(topk, unit=" k value"):
        col_name = f"k_{k:0.0f}"
        scan_name = f"scan_k_{k:0.0f}"
        df = df.groupby(["scan", "window_index"]).head(k)
        prec = df.groupby("match_precursor").head(1).copy(deep=True)
        prec = (prec.groupby("group")
                    .apply(_estimate_qvalues, desc=desc, plus_one=plus_one,
                           col_name=col_name))

        qvals = prec.set_index("precursor")[["scan", col_name]] \
                    .loc[prec.target.values] \
                    .rename(columns={"scan": scan_name})

        results.append(qvals)

    del df # get back some memory.
    return pd.concat(results, axis=1)


# Private Functions -----------------------------------------------------------
def _parse_matrix(psm_matrix: Union[str, pd.DataFrame], desc: bool,
                  window_index: int) -> pd.DataFrame:
    """
    Parse the matrix and transform it for FDR estimation

    This function transforms the wide matrix into a long dataframe,
    and removes missing values.

    Parameters
    ----------
    psm_matrix: str or pandas.DataFrame
        Parse the file containing the matrix or use the provided
        pandas.DataFrame

    desc: bool
        Are higher scores better?

    window_index : int
        The order of this matrix in the input. This is a precaution
        in case the multiple windows share the same scan numbers,
        such as in the case they originated from multiple raw files.
    """
    if isinstance(psm_matrix, str):
        if psm_matrix.endswith(".txt"):
            mat = pd.read_csv(psm_matrix, sep="\t", index_col=0)
        elif psm_matrix.endswith(".pkl.gz"):
            mat = pd.read_pickle(psm_matrix)
        else:
            raise ValueError("Unrecognized file type.")
    else:
        mat = psm_matrix

    mat = (mat.reset_index()
              .rename(columns={"index": "precursor"})
              .melt(id_vars="precursor", value_name="score", var_name="scan")
              .dropna()
              .sample(frac=1)
              .reset_index(drop=True)
              .sort_values(by="score", ascending=(not desc)))

    cols = ["file_idx", "target", "sequence", "charge", "match_sequence"]
    key_df = mat.precursor.str.split("_", expand=True)
    key_df.columns = cols

    mat = pd.concat([mat, key_df], axis=1)
    del key_df

    mat["key"] = mat.precursor
    mat["match_precursor"] = mat["match_sequence"] + "_" + mat["charge"]
    mat["precursor"] = (mat["file_idx"] + "_" + mat["sequence"]
                        + "_" + mat["charge"])

    mat["window_index"] = window_index
    mat["target"] = mat["target"].replace({"T": True, "D": False})

    return mat


def _estimate_qvalues(df: pd.DataFrame, plus_one: bool = False,
                      desc: bool = True, col_name="q-values") \
        -> pd.DataFrame:
    """
    Estimate q-values using the target decoy approach.

    The FDR is estimate simply as the number of targets / decoys at each
    score threshold. Alternatively, a more conservative estimate can be
    obtained by setting 'plus_one=True', which adds one to the numerator.
    The function returns q-values, which are the minimum FDR at which
    the PSM is accepted.

    Parameters
    ----------
    metric : numpy.ndarray
        A 1D array containing the score by which to rank the PSMs.

    target : numpy.ndarray
        A 1D array indicating if the entry is from a target or decoy
        hit. This should be boolean, where `True` indicates a target
        and `False` indicates a decoy. `target[i]` is the label for
        `metric[i]`; thus `target` and `metric` should be of
        equal length.

    desc : bool
        Are higher scores better? `True` indicates that they are,
        `False` indicates that they are not.

    col_name : str
        The name of the new column. The default is "q-values".

    Returns
    -------
    pandas.DataFrame
        A DataFrame with a new column containing the estimated q-value
        for each entry.
    """
    metric = df.score.values
    target = df.target.values

    msg = "'{}' must be a 1D numpy.ndarray or pandas.Series"
    if not isinstance(metric, np.ndarray) or len(metric.shape) != 1:
        raise ValueError(msg.format("metric"))

    if not isinstance(target, np.ndarray) or len(target.shape) != 1:
        raise ValueError(msg.format("target"))

    if not isinstance(desc, bool):
        raise ValueError("'desc' must be boolean (True or False)")

    if metric.shape[0] != target.shape[0]:
        raise ValueError("'metric' and 'target' must be the same length")

    try:
        target = np.array(target, dtype=bool)
    except ValueError:
        raise ValueError("'target' should be boolean.")

    # Sort and estimate FDR
    if desc:
        srt_idx = np.argsort(-metric)
    else:
        srt_idx = np.argsort(metric)

    metric = metric[srt_idx]
    target = target[srt_idx]
    cum_targets = target.cumsum()
    cum_decoys = ((target-1)**2).cumsum()
    num_total = cum_targets + cum_decoys

    # Handles zeros in denominator
    if plus_one:
        cum_decoys += 1

    fdr = np.divide(cum_decoys, cum_targets,
                    out=np.ones_like(cum_targets, dtype=float),
                    where=(cum_targets != 0))

    # Calculate q-values
    unique_metric, indices = np.unique(metric, return_counts=True)

    # Some arrays need to be flipped so that we can loop through from
    # worse to best score.
    fdr = np.flip(fdr)
    num_total = np.flip(num_total)
    if not desc:
        unique_metric = np.flip(unique_metric)
        indices = np.flip(indices)

    qvals = _fdr2qvalue(fdr, num_total, unique_metric, indices)
    qvals = np.flip(qvals)
    qvals = qvals[np.argsort(srt_idx)]

    df[col_name] = qvals
    return df


@nb.njit
def _fdr2qvalue(fdr, num_total, met, indices):
    """
    Quickly turn a list of FDRs to q-values

    Parameters
    ----------
    fdr : numpy.ndarray
        The estimated FDR of each PSM.

    num_total : numpy.ndarray
        The cumulative number of PSMs at each point in the list.

    met : numpy.ndarray
        The unique values of the metric.

    indices : tuple of numpy.ndarray
        The PSMs that have share the same metric value.

    Returns
    -------
    np.ndarray
        The q-value for each PSM. This array will be the same length as
        the provided 'fdr' array.
    """
    min_q = 1
    qvals = np.ones(len(fdr))
    group_fdr = np.ones(len(fdr))
    prev_idx = 0
    for idx in range(met.shape[0]):
        next_idx = prev_idx + indices[idx]
        group = slice(prev_idx, next_idx)
        prev_idx = next_idx

        fdr_group = fdr[group]
        n_group = num_total[group]
        curr_fdr = fdr_group[np.argmax(n_group)]
        if curr_fdr < min_q:
            min_q = curr_fdr

        group_fdr[group] = curr_fdr
        qvals[group] = min_q

    return qvals


# MAIN ------------------------------------------------------------------------
def main():
    """This is executed when run as a command line program"""
    st_time = time.time()
    logging.basicConfig(level=logging.INFO, format="%(levelname)s: %(message)s")

    desc = """
    Estimate q-values using the top-k matches per spectrum.
    When k = 1, this is equivalent to the spectrum-centric search. When
    k = inf, this is equivalent peptide-centric search. Practically, the
    maximum value of k will not exceed the value used for the 'top-match'
    parameter during the tide-search run.
    The format of the 'psm_matrix'---which can be provided either as the
    a file name or a pandas dataframe---should be such that rows are
    precursors, columns are scans, and values are the score. Furthermore,
    the precursor index should be of the format <T|D>_<sequence>_<charge>
    because this string is parsed to determine if the precursor was a
    target or decoy hit. Matrices of this type are saved by the
    'precursor_matrix.py' script.
    """

    parser = argparse.ArgumentParser(description=desc)
    parser.add_argument("psm_matrix", type=str, nargs="+",
                        help="One or more files containing a matrix of PSMs.")

    parser.add_argument("--output_file", type=str, default="precursors.txt",
                        help="The name of the output tab-delimited file.")

    parser.add_argument("--topk", type=str, default="1",
                        help=("One or more integers to use for k. "
                              "Multiple integers can be specified as a comma-"
                              "delimited list. Inf is always used."))

    parser.add_argument("--groups", type=str, default=None,
                        help=("A tab-delimited file with two columns to specify "
                              "groups for group FDR. The first column should be "
                              "the precursor, and the second should be the "
                              "group"))

    parser.add_argument("--ascending", default=False, action="store_true",
                        help=("Lower scores are better."))

    parser.add_argument("--plus_one", default=False, action="store_true",
                        help=("Add 1 to the numerator of the FDR calculation "
                              "for a more conservative estimate."))

    parser.add_argument("--overwrite", default=False, action="store_true",
                        help="Overwrite the existing result file.")

    parser.add_argument("--seed", type=int, default=42,
                        help=("The random seed. Ties are randomly broken "
                              "so setting the seed ensures reproducibility."))

    # Parse arguments
    args = parser.parse_args()

    # Set the random seed
    np.random.seed(args.seed)

    # Parse the topK parameter
    topk = [int(i) for i in args.topk.split(",")]

    # Check that output doesn't exist if overwrite is False
    if os.path.isfile(args.output_file) and not args.overwrite:
        raise RuntimeError(f"{args.output_file} already exists. If you want to"
                           " proceed anyway, use '--overwrite'.")

    # Estimate q-values
    res = assign(psm_matrix=args.psm_matrix, topk=topk, plus_one=args.plus_one,
                 desc=(not args.ascending), groups=args.groups)

    # Save file
    res.to_csv(args.output_file, sep="\t")

    # Wrap-up
    end_time = time.time()
    logging.info("Completed in %0.2f min.", (end_time-st_time)/60)

if __name__ == "__main__":
    main()
```
